# Climate at seed origin drives germination and post-germination trait responses to warming in sessile and pubescent oaks

**DOI:** 10.64898/2026.06.24.734244

**Authors:** Marion Carme, Eduardo Vicente, Marta Benito Garzón

## Abstract

Tree early life stages are particularly sensitive to warming, yet their responses remain poorly understood despite their importance for forest regeneration. Here, we investigated how warming affects early-life traits in two widespread European white oaks: *Quercus pubescens* and *Q. petraea*. We conducted a common garden experiment using 17 populations exposed to three temperature regimes. We measured 19 traits encompassing germination, phenology, and functional and fitness-related traits and performed individual trait mixed-effects models based on temperature transfer distance and the climate of the population. We found that population climate was the primary driver of early stages traits responses to warming, with climatic drivers varying strongly among traits and species. Particularly in *Q. pubescens*, warmer and drier populations showed lower fitness (germination and survival percentages, total biomass) that declined further under warming, consistent with a cost of drought avoidance strategies under continuously wet conditions; in *Q. petraea*, continental populations outperformed others at low temperature transfer distance but suffered the steepest fitness declines under further warming, suggesting a narrow thermal optimum shaped by cold adaptation. Warming generally advanced germination and leaf emergence, increased leaf pigment concentrations and fine-root allocation, reduced specific leaf area. Extreme warming reduced survival, growth and germination. Nevertheless, moderate warming (+0 to +5°C) was rarely detrimental and sometimes beneficial. Our results demonstrate that population climatic origin is a key determinant of regeneration responses to warming, highlighting the need to consider within-species adaptive variation to understand forest regeneration potential under climate change.

## 1. Introduction

Climate change is altering temperature and precipitation regimes, and increasing the frequency and intensity of droughts, fires, storms, and pest outbreaks in forests worldwide **(Dale et al., 2001; Seidl et al., 2017)**. As climatic conditions shift, tree populations may experience local extinction, track suitable environments through migration, or persist in situ by adjusting to novel conditions via evolutionary and ecological strategies **(Aitken et al., 2008)**. Despite their central role in regeneration, early life stages remain comparatively understudied along broad environmental gradients, with most research focusing on adult trees **(Alberto et al., 2013; Leites and Benito Garzón, 2023)**. Yet, germination and post-germination stages represent critical demographic bottlenecks that largely determine recruitment success and long-term population viability **(Máliš et al., 2016; Verdú and Traveset, 2005)**, are key for adaptation to changing environments **(Donohue et al., 2010; König et al., 2022)** and are often more sensitive to climatic variation than adult trees, exhibiting higher mortality and stronger stress responses **(Johnson et al., 2011)**.

Temperature accelerates germination phenology by controlling enzymatic activity and metabolism **(Benavides et al., 2015; Moler et al., 2021)**, and tend to favor germination percentage under wet conditions particularly in the case of temperate species **(Vicente and Benito Garzón, 2024)**. Regarding seedling, warming may increase fitness, e.g. biomass (Zhou et al., 2022), but also impose heat stress **(Grossiord et al., 2022; Murphy and Way, 2021; Teskey et al., 2015)**. As germination and post-germination survival provide integrative proxies of early fitness **(Howard and Goldberg, 2001; Younginger et al., 2017)**, understanding the effects of warming on a comprehensive set of germination and post-germination traits remains crucial to understand the potential for forests seed regeneration under climate change.

Early establishment involves trait coordinated responses across multiple functional dimensions. Seed provisioning determines initial resource availability **(Bartlow et al., 2018; Gómez, 2004)**. Germination and first leaf timing influences growing season length and exposure to seasonal stressors **(Donohue, 2005)**. Growth, structural and architecture traits reflect resource-use strategies, with notably high SLA and rapid growth characterizing acquisitive strategies, vs. conservative with low SLA and slower growth **(Pierce et al., 2013; Wigley et al., 2016)**; acquisitive phenotypes also tend to be taller and slenderer, whereas conservative phenotypes invest relatively more in diameter **(Poorter et al., 2006, 2003)**. Root allocation is closely linked to drought adaptation, with increased root-to-shoot ratios enhancing water acquisition under dry conditions and often decreasing under warm, well-watered conditions **(Ledo et al., 2018)**.

Populations differentiated across environmental gradients may respond heterogeneously to warming, with some locally adapted to current conditions that may face maladaptation under climate change **(Aitken et al., 2008; Alberto et al., 2013)**. The climate driving population differentiation differs between species and populations inhabiting different environmental conditions **(Fréjaville et al., 2020; Mátyás, 2021)**, and also can differ among traits **(Ramírez-Valiente et al., 2021; Vicente et al., 2025)**. These different selection pressures make difficult to assess the effects of warming on trait syndromes across species **(An et al., 2017; Lazarus et al., 2018; Losch et al., 2025; Meeussen et al., 2022)**.

Oaks are foundation forest tree species that play a key role in ecosystem structure and functioning. Among them, *Quercus pubescens* (pubescent oak) and *Q. petraea* (sessile oak) are widespread European white oaks spanning contrasting climatic gradients, and potentially highly vulnerable to climate change **(Baskin and Baskin, 2022; Pritchard et al., 2022; Wyse et al., 2018)**. *Q. pubescens* inhabits Mediterranean-to-temperate climates and exhibits drought-adapted, conservative strategies **(And and Rambal, 1995; Pasta et al., 2016)**. *Q. petraea* inhabits temperate climates and generally shows more acquisitive strategies **(Eaton et al., 2016; Rodríguez-Calcerrada et al., 2008)**. Both typically disperse and start to germinate in autumn/early winter, with a majority of germination and seedling emergence completed in spring **(Joët et al., 2016; Kunstler et al., 2004; Shaw, 1968)**, often later for *Q. petraea* to minimize frost damage risk **(Černý et al., 2024)**. Their recalcitrant seeds (non-dormant, desiccation-sensitive) make establishment success entirely dependent on current-year environmental conditions **(Berjak and Pammenter, 2008)**. Although warming can increase shoot height growth, and reduced stem diameter, root length and biomass in both species **(Arend et al., 2011; Wellstein and Cianfaglione, 2014)**, the magnitude and direction of these responses often vary among populations, as shown in similar traits measured in sapling in adult trees **(Arend et al., 2011; Girard et al., 2022; Jensen and Hansen, 2008; Losch et al., 2025; Mátyás, 2021; Sáenz-Romero et al., 2017; Wilkinson et al., 2017)**.

Here, we combined a common garden experiment with three warming treatments and optimal water availability under controlled conditions, with generalized linear mixed-effects models to investigate how warming and population climate jointly shape early-stage trait responses in *Q. pubescens* and *Q. petraea*. We measured 19 traits spanning fitness (acorn mass, germination, seedling survival and biomass) phenology (germination and first leaf time), resources-use strategy (chlorophyll and leaf photoprotective pigments, SLA, height growth, diameter, height), and rooting (root to shoot and fine to main roots ratios), across 17 range-wide populations. We addressed three main questions: (i) is early-stages trait variation mainly explained by population climate, plasticity to warming, or their interaction?; (ii) how do early-stages traits and their plasticity to warming vary depending on the population climate of origin (i.e. genetic differentiation)?; (iii) how does experimental warming generally affect early-stages trait responses along large environmental gradients? Together, these questions allow us to reveal how adaptation strategies can generate contrasting vulnerabilities to warming at the seedling stage, with important implications for assessing forest regeneration potential of recalcitrant-seeded species under climate change.

## 2. Material and Methods

### 2.1. Plant material

We collected acorns from respectively 7 and 8 natural populations distributed across *Q. pubescens* and *Q. petraea* distribution ranges (**Figure 1**) during autumns 2023 and 2024. Population was considered as a group of at least 10 mature individuals within a single forest stand and thus experienced the same local environmental conditions. We sampled 11 mother trees on average in each population, upon availability (108 mother trees in total) and collected about 25 acorns per mother tree. This sample size was suitable for capturing population responses while incorporating within-population variability **(Mérian et al., 2013)**, particularly given the generally high germination rates reported for *Quercus* species **(Bonner and Karrfalt, 2008)**. The geographical coordinates of the populations and collection date were recorded for all sampled trees (**Figure 1**). Before sowing, we imbibed the acorns in water for 10 minutes and removed the floating ones to sort out the healthy acorns. In total, we used 2112 acorns.

**Figure 1:**
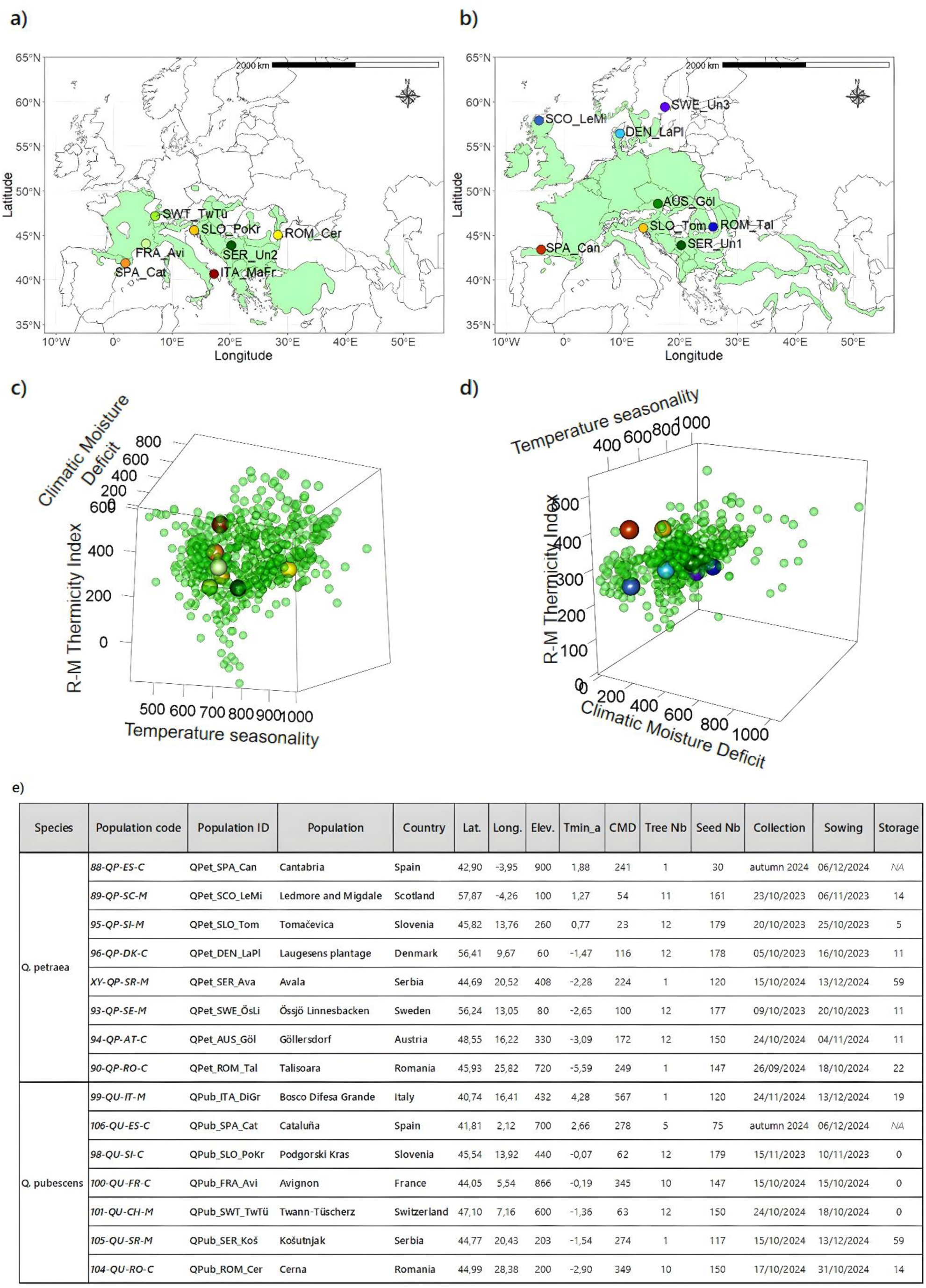
Q. pubescens (on the left) and Q. petraea (on the right) oaks acorn populations within the species distribution range (EUFORGEN, (Caudullo et al., 2017) in a) and b), within the species’ climatic envelope according to Climatic Moisture Index, Rivas-Martínez Thermicity Index and Temperature Seasonality in c) and d). Details about population are in e) Population code is the OptFORESTS code (European project that allowed seed collection in each population); Population ID is the one used in this paper; Tmin_a = population minimal annual temperature; CMD = population Climatic Moisture Deficit; Elev. = Elevation (m); Tree Nb = number of trees per population; Seed Nb = number of acorns per population; Collection = collection date; Sowing = sowing date; Storage = duration of seed storage between collection and sowing (days). Populations colors indicate their origin temperatures (warm colors for warmer populations and cold colors for colder populations, according to mean spring temperature from 1901 to 1960).

### 2.2. Germination experimental design

Acorns were sown as soon as they were collected to prevent desiccation. Hence, the first experiment started on October 2023, and lasted until the end of June 2024, and the second experiment started on November 2024 and lasted until the end of July 2025. Acorns were put in nursery trays (4×5 pots), in 6×6×8 cm pots (one acorn per pot). *Q. petraea* acorns were sown in a standard substrate (NPK 8-2-7; 80% dry organic matter; conductivity 30 mS/m; pH 6.5; water retention 820 mL/L). For *Q. pubescens*, we added perlite at a ratio of two-thirds standard substrate to one-third perlite to increase substrate porosity and drainage and approximates the dry, well-drained soil conditions typical of *Q. pubescens* habitats **(Pasta et al., 2016)**. The apical end of acorns was exposed at the soil surface, allowing direct observation of radicle protrusion.

To test the temperature effect on germination, we sowed the acorns in three climatic chambers (Snijder LABS, micro clima-series) at 15, 20 and 25 °C during the day, and 10, 15 and 20 °C during the night. Other environmental parameters were kept constant: 75% relative air humidity, 100 μmol.m^-2^.s^-1^ light intensity (Li-190R Quantum Sensor), photoperiod of 13/11 light/dark hours, and watering thrice a week with distilled water.

### 2.3. Phenotypic measurements

We measured a comprehensive set of traits to characterize early stages of trees, from acorn mass to seedling survival (**Table 1a**). We measured individual acorn mass (AM) (precision of 10^−1^ g) and fresh and dry population acorn mass (2 acorns per mother tree, acorns dried at 60°C during one week) to calculate population moisture content (MC). Germination (G) and first leaf emergence (FL) were then monitored thrice a week in the three climatic chambers until emergence (G and FL = 1 if germinated/first leaf, 0 if not). An acorn was considered as germinated when radicle had emerged by 2 mm. Germination time (GT) and first leaf time (FLT) were considered as the number of days between sowing and emergence. Height at first leaf emergence (HFL) was also measured. Seedling traits were measured one month after first leaf emergence: survival (S = 1 if alive, 0 if not); height (H); diameter (D); root collar (RC); leaves pigments: chlorophyll (Chl), flavonols (Flv), anthocyanins (Anth), nitrogen-flavonols index (NFI) using an MPC-100 multi-pigment meter (ADC BioScientific Ltd, UK); leaves number, leaf surface and mass, main root mass, fine roots mass, stem number, main root number and aboveground biomass.

**Table 1:** Phenotypic trait measurements in a) and indexes calculated from measurements in b).

| Trait type | Trait | Measurement detail and unit |
| --- | --- | --- |
| Seed | Fresh acorn mass | gram |
|  | Moisture content | Averaged moisture content measured for each population. Acorns were dried more than 1 week at 60°C. (percentage) |
| Germination phenology | Germination time | Number of days between sowing and germination emergence (days) |
| Germination | Germination (status) | An acorn was considered as germinated when radicle had emerged by 2 mm. Monitored every 2 – 3 days. G = 1 if germinated, 0 if not. |
| Seedling phenology | First leaf emergence time | Number of days between sowing and first leaf emergence (days) |
| Seedling traits | Height at first leaf emergence | Centimeters. |
|  | Survival (status) | S = 1 if alive, 0 if not. Measured one month after first leaf emergence. |
|  | Height | Stem height (centimeters). Measured one month after first leaf emergence. |
|  | Diameter | Stem diameter at the base (millimeters). Measured one month after first leaf emergence. |
|  | Root collar | Root collar at the base (millimeters). Measured one month after first leaf emergence. |
|  | Leaves number | Measured one month after first leaf emergence. |
|  | Leaf surface | One leaf surface (cm<sup>2</sup>), measured with Petiole Pro application. Measured one month after first leaf emergence. |
|  | Leaf mass | The same leaf dry mass (gram). Measured one month after first leaf emergence. |
|  | Leaves pigments | Chlorophyll, flavonoids, anthocyanins, nitrogen-flavonoids index (NFI). Measured using an MPC-100 multi-pigment meter (ADC BioScientific Ltd, UK), one month after first leaf emergence. |
|  | Main root(s) mass | Dry mass of main root(s) (gram). Measured one month after first leaf emergence. |
|  | Fine roots mass | Dry mass of fine roots (gram). Measured one month after first leaf emergence. |
|  | Aboveground biomass | Dry mass of stem and leaves (gram). Measured one month after first leaf emergence. |
|  | Stem number | Number of stems. Measured one month after first leaf emergence. |
|  | Main root number | Number of main roots. Measured one month after first leaf emergence. |
|  | Belowground biomass | Dry mass of main and fine roots |

| Index | Formula | Formula explanation |
| --- | --- | --- |
| Relative vertical growth rate (consider the fact seedling size) | $RGR = \frac{\ln(H) - \ln(HFL)}{30}$ | RGR is the vertical Relative Growth Rate; H is the seedling height one month after leaf emergence; HFL is the seedling height at first |
| vary initially at first leaf emergence) |  | leaf emergence; 30 is the number of days between the 2 heights measurements |
| Specific Leaf Area | $SLA = \frac{LS}{LM}$ | SLA is the Specific Leaf Area index; LS is the surface of one leaf (cm<sup>2</sup>); LM is the dry mass of this same leaf |
| Root to shoot ratio | $RtS = \frac{UGB}{AGB}$ | RtS is the root to shoot ratio; UGB is underground biomass (the sum of fine roots and main root mass); AGB is aboveground biomass (the sum of stem and leaves biomass) |
| Fine root to main root ratio | $FtMR = \frac{FRM}{MRM}$ | FtMR is fine root to main root ratio; FRM is fine roots mass; MRM is main root mass. |
| Total biomass | $B = UGB + AGB$ | UGB is underground biomass; AGB is aboveground biomass |

From these measurements, we calculated several supplementary indexes for each combination of population – chamber – mother tree (**Table 1b**).

### 2.4. Climatic data

To characterize the climate of origin for each population, we retrieved 28 climatic variables from ClimateDT over the 1901-2023 period **(Marchi et al., 2024)**. Climatic conditions were summarized over two temporal windows reflecting different, but not mutually exclusive, dimensions of population climate exposure: (i) a long-term climatic window (1901–1960), representing historical climate conditions under which populations likely differentiated (here, the average climate from 1901 to 1960 i.e the beginning of available data to the proxy of the current climate change onset); (ii) a short-term window (the year preceding the acorn collection, in 2023 and 2024), representing climatic conditions during acorn development **(Olson and Boyce, 1971)**.

From these, we selected a subset of non-correlated variables that captured key climatic dimensions (water availability, temperature, and seasonality) for each population (**Figure S1**; **Table 2**).

**Table 2:** Population climatic variables selected to characterize populations and investigate population effect on germination and seedling traits in generalized linear models.

| Name and Acronym | Unit | Description | Calculation |
| --- | --- | --- | --- |
| Average monthly maximal temperatures for each season (Tsp, Tsm, Taut, Twt) | °C | Population temperature |  |
| Isothermality (IsoT) | ratio | Quantifies the relative amplitude of daily temperature fluctuations compared to annual temperature fluctuations | Temperature Diurnal Range / Temperature Annual Range |
| Temperature seasonality | °C | Annual temperature fluctuations | Temperatures standard deviation x 100 |
| Minimal annual temperature (Tmin_a) | °C | Coldest population temperature, exposure to extreme cold events |  |
| Average montly precipitations for each season (Psp, Psm, Paut, Pwt) | mm | Population water input |  |
| Hargreaves Climatic Moisture Deficit (CMD) | mm | Population water balance, drought stress degree | Hargreaves reference evapotranspiration (computed with the SPEI R package) minus mean annual precipitation (Hargreaves and Samani, 1985) |

To characterize the experiment temperature related to population climate, we used a temperature transfer distance approach. We calculated the temperature transfer distance (TD, °C) as the difference between the climatic chamber temperature and each population spring temperature, and is calculated as follows:

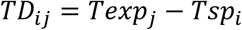

where TD*_ij_* is the temperature transfer distance of the growing season for population *i* in chamber *j*; Te*x*p is the day temperature of the climatic chamber *j*; T*s*p is the mean spring temperature of the population *i* for the period 1900-1960.

### 2.5. Statistical analysis

All analyses were carried out in R version 4.3.2 **(R Core Team, 2023)** and script is available upon publication.

#### 2.5.1. Generalized linear mixed-models of the effects of population and sowing temperature on early stages

For each trait and each species, we built a generalized linear mixed-effects model to test for the effects of the seeds’ climate of origin, temperature transfer distance and its interaction. They took the following general form:

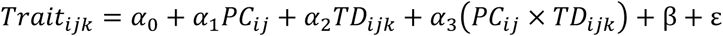

where i*jk* stands for the *i* th acorn of the *j* th population at the *k* th temperature. *TD* is the temperature transfer distance; *PC* are the population climate variables; *α* are the coefficients to be estimated; *β* is the mother tree nested in the population as random effect; *ε* the residuals.

To select explanatory variables, we performed multiple variable selection methods for each model (**Table S1**). We then performed multiple model comparisons via ANOVA and Akaike Information Criterion (AIC), to test for the effects of these variables, their quadratic terms, and their interactions. We also tested the effect of storage, which was generally not significant.

The models for each species and trait are described in supplementary material (**Table S2**). The choice of the model type depended on the statistical distribution of each trait. For acorn mass, we built gamma regression models using a log link function (function *glmmTMB* from the glmmTMB 1.1.8 R package **(Brooks et al., 2017)**). For germination, survival, and stem number, we built logistic mixed-effects models using a logit link function (function *glmer* from the lme 4 1.1-35.1 R package **(Bates et al., 2015)**). For germination time and first leaf emergence time, we built generalized Poisson mixed-effects model using a log link function (function *glmmTMB* from the glmmTMB 1.1.8 R package **(Brooks et al., 2017)**). For relative growth percentage, we built beta regression models using a logit link function (function *glmmTMB* from the glmmTMB 1.1.8 R package **(Brooks et al., 2017)**). For total biomass, specific leaf area, chlorophyll, flavonols, NFI, anthocyanins, height, diameter, root collar, root to shoot ratio and fine to main root ratio, we built Gaussian regression models using an identity link function (function *glmmTMB* from the glmmTMB 1.1.8 R package **(Brooks et al., 2017)**).

#### 2.5.2. Models evaluation and variance partitioning

The variance explained by the GLMMs of the effects of population and sowing temperature on germination was assessed using different R² metrics: marginal (R2M) and conditional (R2C) for all mixed-effects models, and the ordinary coefficient of determination R² for beta regression models. These calculations were performed using the *model_performance* function from performance 0.10.8 R package **(Lüdecke et al., 2021)**. Residuals plots and Variance Inflation Factors were used to validate the suitability of the models. Tables and plots related to model evaluation are available in Supporting Information (**Table S3**). All explanatory variables were standardized before running the models. We assessed the unique contributions of predictors i.e. variance explained, using partR2 0.9.2 R package **(Stoffel et al., 2021)** (**Table S4**; **Figure S1**).

## 3. Results

### 3.1. Relative contributions of population climate and temperature transfer distance to early life-stage traits

Population climate was the most important variable explaining germination time and percentage, survival, total biomass, root to shoot ratio, and flavonols in both species. Furthermore, climate of the population was the most important driver of height, diameter and root collar variation in *Q. pubescens*, and of first leaf emergence time, specific leaf area and NFI in *Q. petraea* (**Figure 2**; **Figure S1**). The population temperature-related variables mostly explained survival, total biomass, diameter, root collar, specific leaf area, whereas water related-variables mostly explained acorn mass and flavonols responses to warming (**Figure S1**).

**Figure 2:**
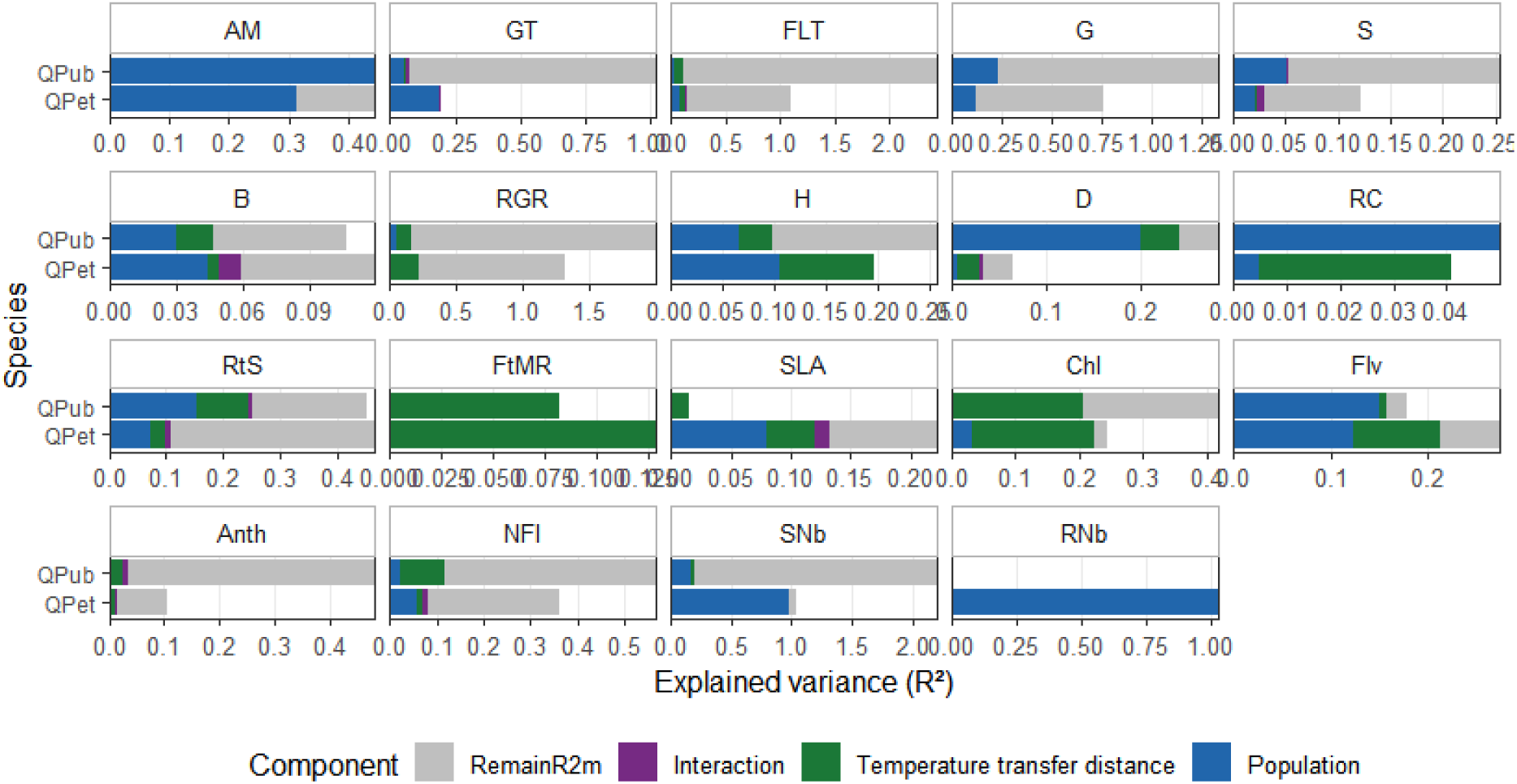
Explained variance partitioning for each generalized linear mixed-model of the effects of population and temperature transfer distance (including experiment temperature) on seed germination and seedling traits in Q. pubescens and Q. petraea. In blue, population climate variables explained part; in green, temperature transfer distance capturing phenotypic plasticity and genetic variation in plasticity; in purple, the interaction between transfer distance and population climate; in grey, the remaining part of marginal R-squared (partR2 quantifies semi-partial r-squared, representing the variance uniquely explained by each predictor; their sum is lower than the model’s marginal R-squared because correlated predictors share explained variance that cannot be uniquely assigned to individual variables). The remaining part of conditional R-squared is not shown for the sake of readability. Traits acronyms mean: AM = Acorn Mass; GT = Germination Time; FLT = First Leaf Emergence time; G = Germination; S = Survival; B = total Biomass; RGR = Relative Growth Rate; H = Height; D = Diameter; RC = Root Collar; RtS = Root to Shoot ratio; FtMR = Fine to Main Roots ratio; SLA = Specific Leaf Area; Chl = Chlorophyll; Flv = Flavonoids; Anth = Anthocynanins; NFI = Nitrogen-Flavonoids Index; SNb = Stem Number; RNb = Root Number.

Fine-to-main root ratio and chlorophyll were mostly driven by the temperature transfer distance. The interactions between population climate and temperature transfer distance explained more variance in *Q. petraea* than in *Q. pubescens* in SLA, total biomass and survival.

For the case of anthocyanins, germination time and percentage, first leaf emergence time, survival percentage, relative growth percentage, and NFI a large fraction of explained variance is shared between them; thus respective total contributions can’t be disentangled in these traits (**Figure 2**).

### 3.2. Fitness-related traits

#### 3.2.1. Acorn mass

Acorn mass model was driven by the precipitation seasonality of the seed origin in *Q. pubescens*. In *Q. petraea*, higher autumn precipitation of the seed origin during the maturation year was the main driver of acorn mass (**Table S4**; **Table S5**; **Figure S3**). The overall performance of acorn mass models was high (**Table S3**).

#### 3.2.2. Germination percentage

The main drivers explaining germination percentage in *Q. pubescens* were the spring temperature of the population (15% of variance explained, main predictor) along with the temperature transfer distance (TD) (**Table S4**): TD decreased germination after an optimum, especially in populations originating from warmer springs, which also were the populations germinating the least (**Figure 3**; **Table S5**). For *Q. petraea* the seasonal population precipitation and the isothermality of the acorn maturation year were the most important drivers together with the TD (**Table S4**): populations from higher isothermality origins germinated less, and TD decreased germination (**Figure 3**; **Table S5**).

**Figure 3:**
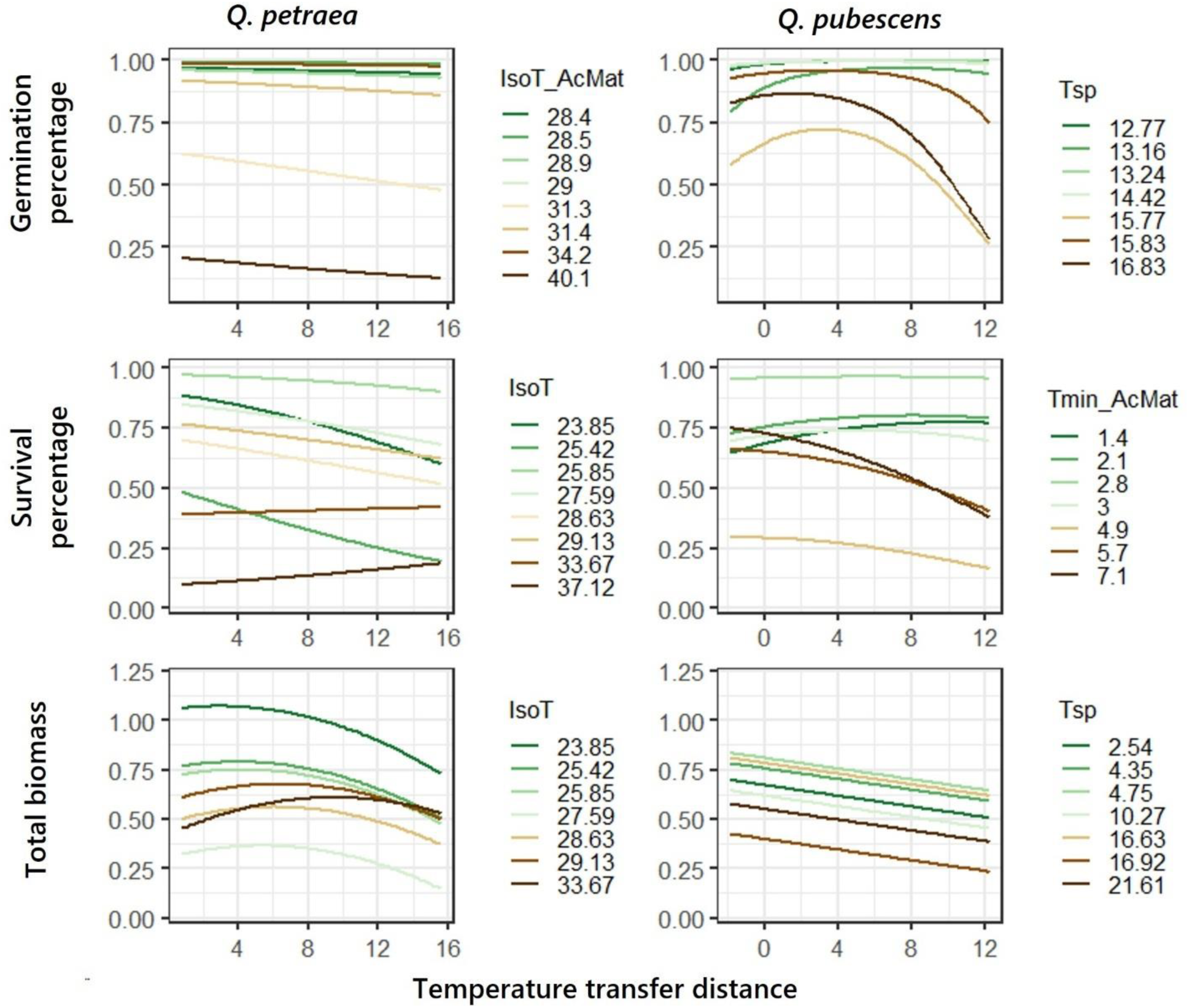
Plots showing the effects of temperature transfer distance and of the most important population climate variable for each generalized linear mixed-model, for germination, survival percentage, and total biomass. Colored lines are populations represented function of the population climate variable that explained most variance. Climate variables acronyms mean: IsoT = Isothermality; Tmin = minimal annual Temperature. No suffix indicates the variable is averaged on the 1901-1960 period. _AcMat suffix indicates the variable is averaged on the acorn maturation year.

#### 3.2.3. Survival percentage and total biomass

The most important population drivers of survival along with TD were isothermality along with TD for both species, and the minimum temperature at the acorn maturation year for *Q. pubescens*. The most important population driver of biomass was isothermality for *Q. petraea*, and spring temperature for *Q. pubescens* (**Table S4**).

Temperature transfer distance often did not have a clear effect on survival and total biomass when moderate (0 - +5°C) and tended to decrease them after an optimum which was not captured in every trait and species. In *Q. pubescens*, this decrease in survival was the strongest in populations originating from warmer spring and winter climates (which were also the populations showing the lowest survival and biomass). In *Q. petraea*, decrease in survival and biomass was the strongest in populations originating from lower isothermality origins (which were also the populations showing the lowest survival and biomass) (**Figure 3**; **Table S5**).

The variance explained by the survival and biomass models for both species was low (marginal R squared of 6, 10% and 8 and 7% respectively) (**Table S3**).

### 3.3. Phenology-related traits

#### 3.3.1. Germination time

The significant drivers of *Q. pubescens* germination time were spring temperature, autumn precipitation along with TD. The driest autumn populations showed the longer germination time, but were also the only populations in which TD decreased germination time (**Figure 4**; **Table S5**).

**Figure 4:**
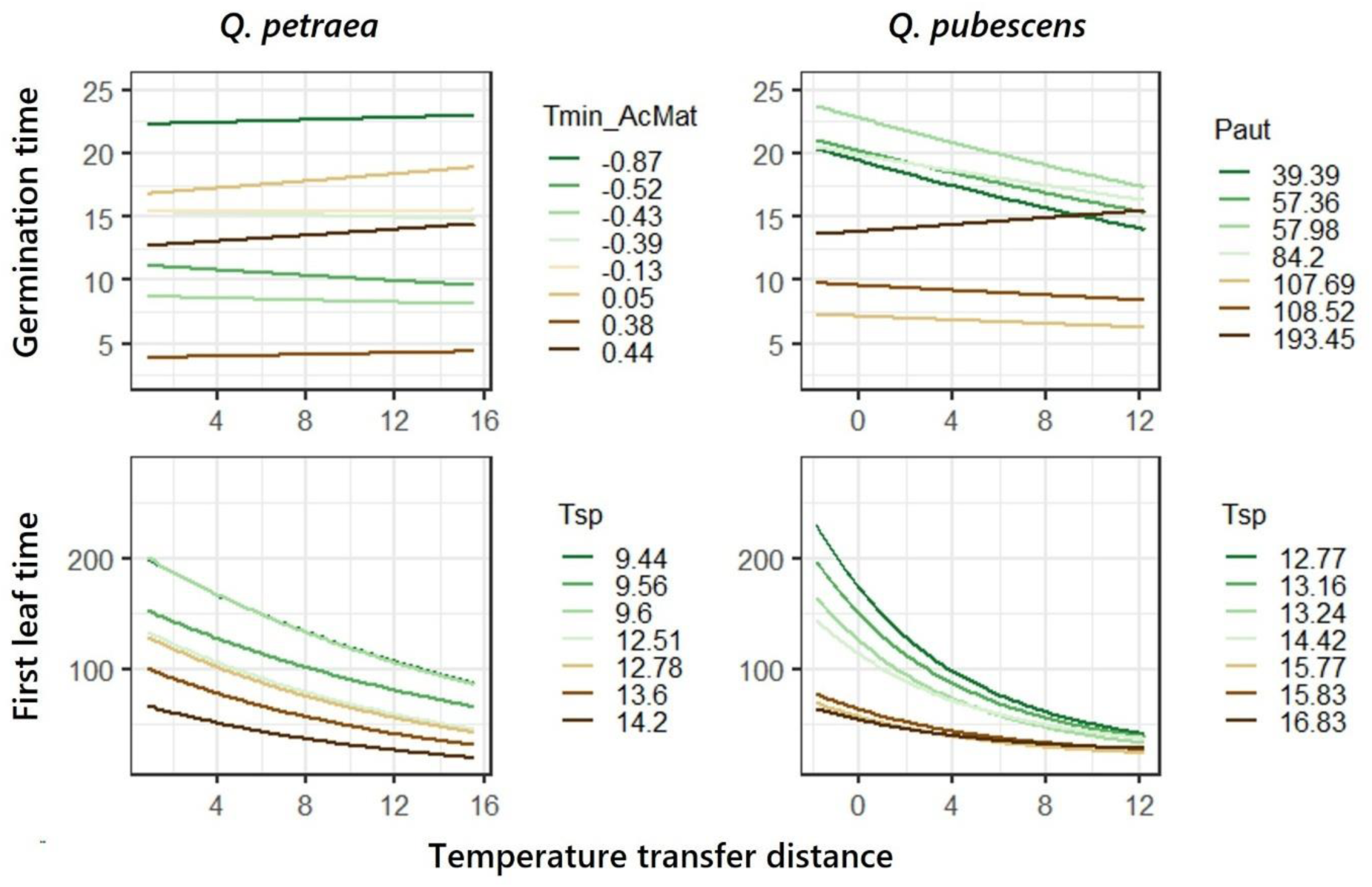
Plots showing the effects of temperature transfer distance and of the most important population climate variable for each generalized linear mixed-model, for germination time and first leaf emergence time. Colored lines are populations represented function of the population climate variable that explained most variance. Climate variables acronyms mean: Tmin = minimal annual Temperature; Tsp = mean spring Temperature; Paut = mean autumn precipitation. No suffix indicates the variable is averaged on the 1901-1960 period. _AcMat suffix indicates the variable is averaged on the acorn maturation year.

The drivers of germination timing in *Q. petraea* were temperature transfer distance together with the minimal temperature of acorn maturation year and the population seasonal precipitation. Population climate explained a lot more variance (15%) than transfer distance (0.3%). Germination time was smaller in populations with warmer winters (**Figure 4**).

#### 3.3.2. First leaf emergence time

The most important population drivers of first leaf emergence were the spring temperature and autumn precipitation for *Q. pubescens* and the annual minimal temperature and precipitation seasonality of acorn maturation year for *Q. petraea* (**Table S4**). The TD was significant and decreased the time to germination especially in warmer spring populations in both species (**Table S5**; **Figure 4**).

### 3.4. Resources-use strategies related traits

#### 3.4.1. Leaf pigments: chlorophyll, flavonols, anthocyanins

Chlorophyll content was especially driven by the transfer distance in *Q. petraea* and *Q. pubescens* (respectively 13% and 25% of variance explained). Flavonol content was more explained by population climate; climatic moisture deficit for *Q. petraea* and spring precipitation for *Q. pubescens* (respectively 12% and 15% of variance explained, with transfer distance reaching 3% maximum).

Anthocyanins leaf content was little explained by both the transfer distance and the climatic moisture deficit for *Q. petraea* and by the transfer distance and the populations’ spring temperature for *Q. pubescens* (**Table S4**).

Higher TD increased all leaf pigment contents (chlorophyll, flavonols, and anthocyanins), with stronger increases in *Q. pubescens* populations from warmer spring climates and in *Q. petraea* populations from sites with higher climatic moisture deficit. Quadratic relationships with temperature were common across pigment traits, including an anthocyanin optimum at approximately 22°C in *Q. pubescens*. Chlorophyll also reached an optimum in both species (around + 15°C for *Q. petraea*, and +10°C in *Q. pubescens*).

*Q. petraea* seedlings coming from populations with high climatic moisture deficit displayed high chlorophyll, anthocyanins, and low flavonols content. In *Q. pubescens*, populations coming from higher spring precipitation displayed high flavonols content. *Q. pubescens* seedlings coming from warmer spring populations displayed high chlorophyll and anthocyanins content. (**Figure 5**; **Table S5**).

**Figure 5:**
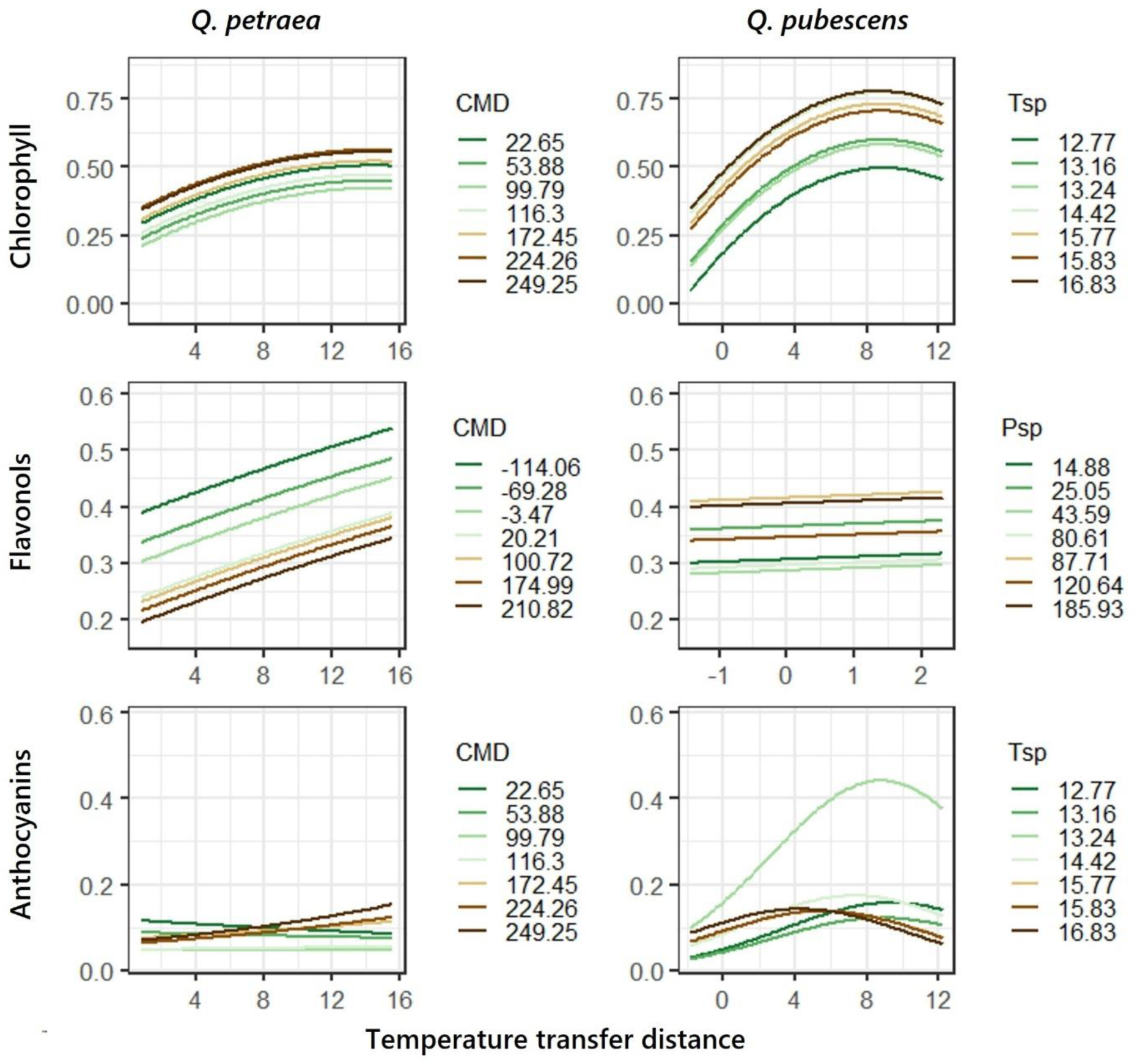
Plots showing the effects of temperature transfer distance and of the most important population climate variable for each generalized linear mixed-model, for chlorophyll, flavonols and anthocyanins. Colored lines are populations represented function of the population climate variable that explained most variance. Climate variables acronyms mean: Tsp = mean spring Temperature; CMD = Climatic Moisture Deficit; Psp = spring precipitation. No suffix indicates the variable is averaged on the 1901-1960 period. _AcMat suffix indicates the variable is averaged on the acorn maturation year.

The variance explained by the anthocyanins *Q. petraea* model was low (marginal R squared of 5 %) (**Table S3).**

#### 3.4.2. Vertical relative growth rate

Relative growth was explained exclusively by the transfer distance in *Q. petraea* whereas in *Q. pubescens* it was explained by the transfer distance and the population spring temperature and precipitation (**Table S4**).

Populations planted at warmer conditions than their origin generally decreased relative growth rate, and in *Q. pubescens,* warmest spring populations always showed the highest growth (**Figure 6**; **Table S5**).

**Figure 6:**
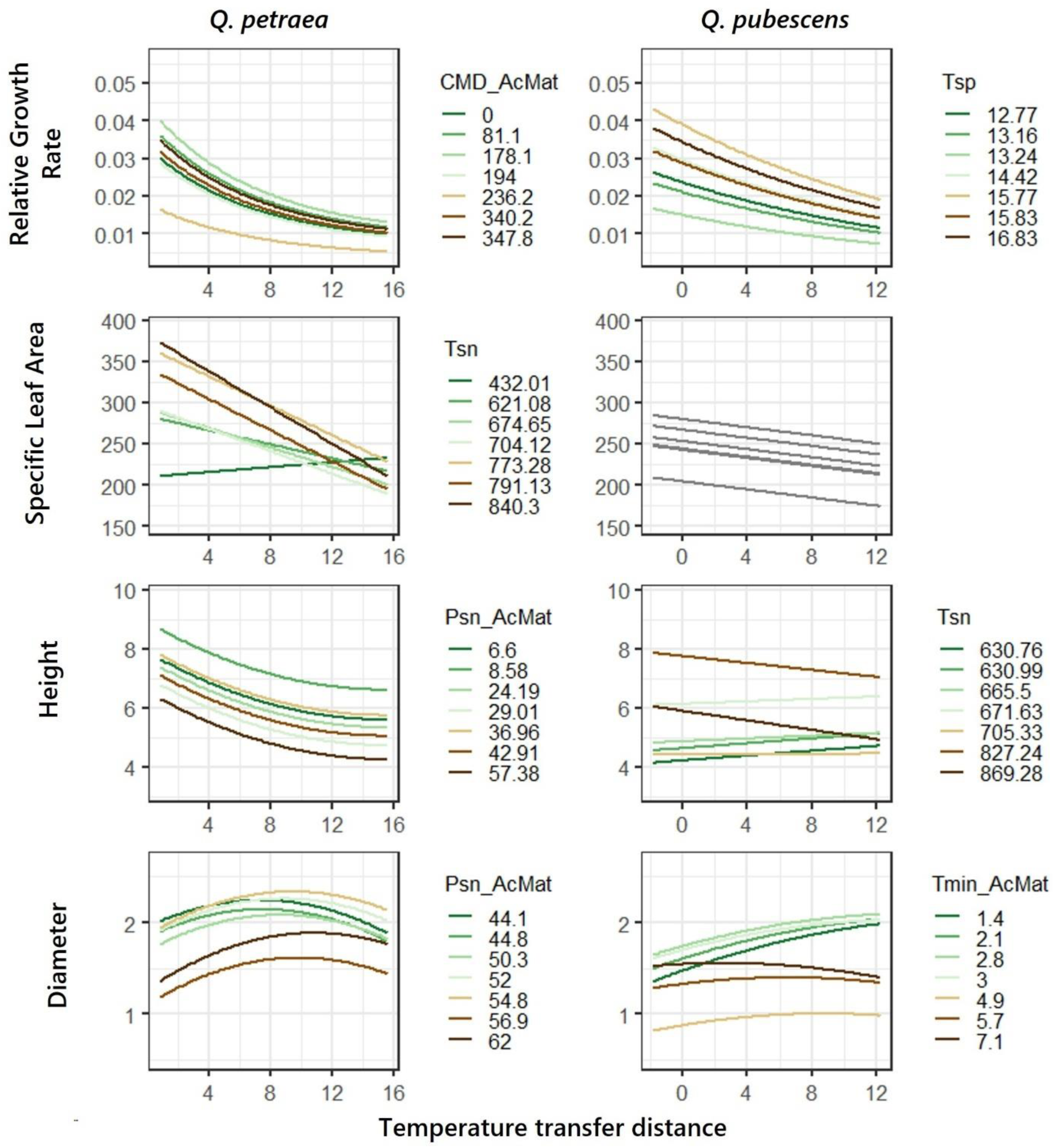
Plots showing the effects of temperature transfer distance and of the most important population climate variable for each generalized linear mixed-model, for relative growth rate, specific leaf area, height and diameter. Colored lines are populations represented function of the population climate variable that explained most variance. Climate variables acronyms mean: Tsp = mean spring Temperature; Tmin = minimal annual Temperature; Psn = Precipitation seasonality; Tsn = Temperature seasonality; IsoT = Isothermality; CMD = Climatic Moisture Deficit. No suffix indicates the variable is averaged on the 1901-1960 period. _AcMat suffix indicates the variable is averaged on the acorn maturation year.

The variance explained by the relative growth models were high (marginal R squared of 77 and 78% for *Q. petraea* and *Q. pubescens*) (**Table S3**).

#### 3.4.3. SLA

SLA main driver was the population temperature seasonality (7% of variance explained) together with the transfer distance (4%) for *Q. petraea*, and the transfer distance alone for *Q. pubescens*.

It tended to decrease with warming in both species (**Figure 6**).

In *Q. petraea*, this response was most pronounced in populations exposed to greater seasonal temperature variation, which also exhibited higher SLA than the other populations (**Figure 6**; **Table S5**).

The variance explained by the SLA *Q. pubescens* model was low (marginal R squared of 1%) (**Table S3).**

#### 3.4.4. Stem height and diameter

The most important variables explaining stem height were the seasonal precipitation (20% of variance explained) and the transfer distance (4%) for *Q. petraea* and equally the transfer distance, the spring temperature and the temperature seasonality for *Q. pubescens* (**Table S4**).

Height decreased under warming in every *Q. petraea* populations and only in populations with higher temperature seasonality in *Q. pubescens* (**Figure 6**; **Table S5**).

The stem diameter was explained by the transfer distance (4% of variance explained) and the seasonal precipitation of the acorn maturation year (3%) for *Q. petraea* and especially by the minimum temperature during the acorn maturation (14%), then spring seasonality and transfer distance for *Q. pubescens* (**Table S4**).

Overall, the stem diameter increased until an optimum when seedlings were grown at warmer than home climates for both species (**Figure 6**). In *Q. petraea*, populations originating from sites with higher precipitation seasonality during the acorn maturation year showed the lowest diameter but also the highest increase in diameter under warming. In *Q. pubescens,* populations from sites with lower minimum temperatures during the maturation year showed the highest increase in diameter under warming (**Figure 6**; **Table S5**).

Stem diameter in *Q. petraea* had low marginal R squared of 4% whereas for the other traits it was higher (**Table S3**).

### 3.5. Rooting-related traits

#### 3.5.1. Root to shoot ratio

The main drivers of the root-to-shoot ratio were the population isothermality (9% of variance explained) and the transfer distance (3%) for *Q. petraea* and the transfer distance (9%) and the population spring temperature (8%) and precipitation seasonality (7%) for *Q. pubescens* (**Table S4**). Seeds planted in warmer-than-home climates displayed smaller root-to-shoot ratios in their seedlings for both species (**Figure 7**). The decrease in root-to-shoot was stronger in *Q. petraea* populations experiencing low isothermality. In *Q. pubescens*, the decrease in root-to-shoot ratio was stronger in populations from warmer spring climates, and populations coming from higher spring temperature areas always displayed smaller root-to-shoot ratio compared to other populations (**Figure 7**; **Table S5**).

**Figure 7:**
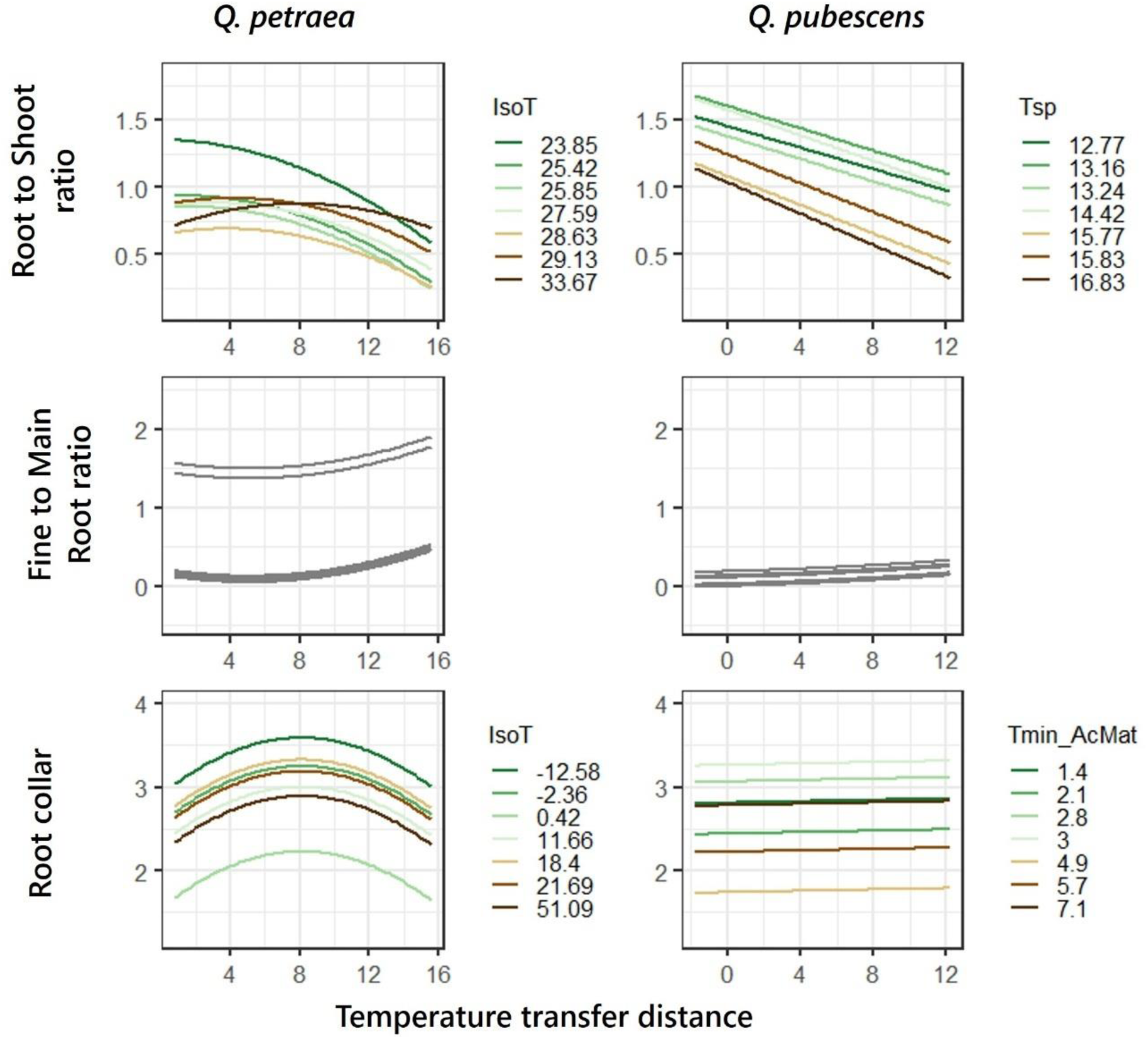
Plots showing the effects of temperature transfer distance and of the most important population climate variable for each generalized linear mixed-model, for root to shoot and fine to main roots ratio and root collar. Climate variables acronyms mean: Tsp = mean spring Temperature; Tmin = minimal annual Temperature; IsoT = Isothermality. No suffix indicates the variable is averaged on the 1901-1960 period. _AcMat suffix indicates the variable is averaged on the acorn maturation year.

Root-to-shoot model showed intermediate variance explained in both species (0.23 and 0.30, respectively) (**Table S3**).

#### 3.5.2. Fine-to-main root ratio

The fine-to-main root ratio was explained by the transfer distance alone in both species (**Table S4**). Seeds planted in warmer-than-home climates displayed higher fine-to-main root ratios in their seedlings for both species (**Figure 7**).

Fine to main root ratio model in *Q. pubescens* had low variance explained (marginal R squared of 8%) (**Table S3**).

#### 3.5.3. Root collar

The main drivers of root collar were the transfer distance and isothermality in *Q. petraea* and the minimum temperature of the acorn maturation year alone for *Q. pubescens* (**Table S4**).

In *Q. petraea,* populations planted in warmer-than-home climates showed an increase in root collar until an optimum at +8°C and then decreased. In *Q. pubescens*, populations coming from low minimum annual temperature displayed bigger root collar diameter (**Figure 7**; **Table S5**).

Root collar models in *Q. petraea* and *Q. pubescens* had low variance explained (marginal R squared of 4%; 5% respectively) (**Table S3**).

## 4. Discussion

### 4.1. Population climate drives early-life trait variation under warming

Although environment effects are generally higher than population effects driving trait variation in saplings and adults in most tree species **(Archambeau et al., 2022; Benito Garzón et al., 2019; Bresson et al., 2011; Liao et al., 2016; Sáenz-Romero et al., 2017; Stamp and Hadfield, 2020)**, our results show that the climate of the population was the main driver of germination percentage, survival, biomass and root:shoot allocation in both species. Thus, there may be a higher influence of the population effect at early stages compared with sapling and adult trees, as it has already been reported for *Abies alba* and *Fagus sylvatica* **(Csilléry et al., 2026; Muffler et al., 2021)**, suggesting that environmental filtering is more important in saplings than right after germination, where the populations effect is crucial for seedling establishment.

The trait responses to warming were driven by very different population variables. For instance, population spring temperature and precipitation were the main drivers of 12/19 and 8/19 traits for *Quercus pubescens* whereas *Q. petraea* traits were mostly driven by the climatic moisture deficit, isothermality and minimum annual temperature in 12/19, 6/19 and 2/19 drivers. These climatic population drivers are expected for a submediterranean and temperate species, which are subject to different selection pressures across their ranges **(Eaton et al., 2016; Pasta et al., 2016)**.

In *Q. pubescens*, warmer and drier populations (higher spring and annual minimum temperatures and lower precipitation) exhibited more acquisitive strategies (higher RGR, lower diameter, lower leaf protective pigments) (**Figure 5**; **Figure 6**) alongside lower baseline fitness (survival, biomass, germination) (**Figure 3**), which declined even further under warming. A similar pattern has been reported in *Q. suber* **(Vicente et al., 2025)**. This seemingly contradicts the expectation that warm-dry origin populations should be best pre-adapted to hotter, drier futures. This can be explained by considering the cost of drought avoidance strategies under the continuously wet conditions of our experiment. In Mediterranean and semi-arid plants, including *Q. pubescens*, strategies where acquisitive behaviour is maintained even under stress to exploit briefly favourable windows have already been documented **(And and Rambal, 1995; Blanco-Sánchez et al., 2022; Nardini and Pitt, 1999; Tognetti et al., 2007; Yan et al., 2025)**, but such pulse-use strategies come at a cost in sustained, non-drought environments **(Cavender-Bares, 2019; Kooyers, 2015; Schwinning and Ehleringer, 2001; Tardieu, 2012)**. This is also consistent with the broader trade-off framework described by **(Willi and Van Buskirk, 2022)**: at the warm-dry distribution edge, rapid growth and direct avoidance strategies are selected, at the expense of performance in other environments.

In *Q. petraea*, more continental populations (lower isothermality) exhibited higher fitness (survival, biomass) under well-watered warm conditions, but experienced the steepest declines under further warming (**Figure 3**), seemingly losing their initial advantage. This runs counter to some range-wide predictions that western, warmer populations would suffer most under future climate change while eastern and northern populations would benefit **(Meeussen et al., 2022; Sáenz-Romero et al., 2017)**, though contrasting predictions also exist **(Mátyás, 2021)**. While our results alone cannot resolve this debate, they help quantify the associated uncertainty. A plausible explanation lies in the asymmetry of thermal adaptation: cold tolerance varies more strongly across latitude than heat tolerance **(Lancaster and Humphreys, 2020)**. Consequently, adaptive differentiation among populations may be greater at the cold than at the warm edge of the thermal niche. Our results are consistent with studies showing that populations adapted to cold climates can exhibit strong negative responses to warming, e.g. in birch and tundra species **(DeMarche et al., 2018; Skre et al., 2017)**, potentially reflecting evolutionary trade-offs whereby some traits values favored under short growing seasons or frequent frost events become less advantageous when temperatures increase **(Willi and Van Buskirk, 2022)**.

### 4.2. Warming responses

#### 4.2.1. Moderate warming does not reduce performance

Germination percentages and post-germination survival and biomass provide integrative proxies of early fitness **(Howard and Goldberg, 2001; Younginger et al., 2017)**, and showed similar responses in our study. Overall, moderate warming (0-5°C) did not obviously reduce fitness, making it or stagnate, or increase until an optimum, or slightly decrease (**Figure 3**), in line with the fact that with enough water and nutrients, moderate warming has positive effects (e.g. warming of 2.5°C increase tree biomass in **(Zhou et al., 2022)** meta-analysis). On the other hand, more extreme warming (+5 to + 16°C) was generally detrimental, indicating strong physiological stress responses even under non-limiting water conditions, as already reported across temperate and subartic species **(Bassow et al., 1994; Ibáñez et al., 2017; Lazarus et al., 2018; Morin et al., 2010; Rubio et al., 2025; Shevtsova et al., 2009; Vicente et al., submitted)**.

Optimum captured above the home climate between +3 and +16°C in germination of both species, survival of *Q. pubescens* and total biomass of *Q. petraea* (**Figure 3**) may reflect an adaptation lag, as it was hypothesized for germination of some Mediterranean, temperate and boreal species **(Vicente and Benito Garzón, 2024).**

#### 4.2.2. Warming hastens first leaf emergence

Seedlings advanced their first leaf emergence when planted at warmer-than-home climate, as expected. Global warming causes advances in early-life phenology especially in juveniles (Sperling et al., 2019; Stuble et al., 2021) for many species, including oaks such as *Q. petraea*, *Q. pagoda*, *Q. pubescens*, and *Q. robur* **(Hawkins, 2019; Jastrzębowski et al., 2021; McCartan et al., 2015)**, by modifying water absorption, liquid diffusion, and enzymatic action in the seed **(Korstian, 1927)**, and affecting the percentage of tissue changes and metabolic processes required for epicotyl emergence and leaf flush **(Hawkins, 2019)**. This advanced phenology in first leaf emergence can be beneficial to avoid summer drought **(Pearson and D’Orangeville, 2022; Warwell and Shaw, 2018)**, although it can be detrimental for populations exposed to late frosts **(Bennie et al., 2010; Gömöry and Paule, 2011; Muffler et al., 2016)**.

Germination timing patterns were generally less consistent and more difficult to interpret. Across species, *Q. petraea* showed little response of germination timing to warming, whereas *Q. pubescens* exhibited population-dependent shifts, with earlier germination under warming primarily in seeds originating from drier climates. This pattern is consistent with the idea that seeds from drier origins may benefit from earlier establishment under warmer conditions, likely increasing the time available for seedling development before the onset of summer stress. However, the ecological meaning of “earlier germination” differs among species depending on their epicotyl dormancy strategies which can decouple radicle emergence from subsequent seedling development **(Baskin and Baskin, 2021; Jaganathan et al., 2025; Joët et al., 2013)**, and should therefore be interpreted with caution.

#### 4.2.3. SLA and leaf pigment content: defense investment and conservative shifts under warming

Most populations increased flavonols and anthocyanin leaf contents under warming in both species, suggesting that warming creates oxidative stress requiring protection even under optimal water conditions. These compounds are related to protection against high temperatures and arid conditions **(Pintó-Marijuan et al., 2013; Shao et al., 2007)**.

Chlorophyll increased under warming in both species up to an optimum, with *Q. pubescens* reaching this optimum at a cooler temperature (+10°C) versus *Q. petraea* (+15°C). The chlorophyll optimum in both species suggests a temperature threshold triggering protective metabolic down-regulation rather than chloroplast damage **(Bielinis et al., 2015; Yüzbaşıoğlu et al., 2017)**. In fact, *Q. pubescens* is able to maintain photosynthetic capacity under heat stress through reversible down-regulation, with chlorophyll reductions reflecting protective responses **(Haldimann et al., 2008; Haldimann and Feller, 2004)**.

SLA decreased under warming in both species, representing a shift toward more conservative leaves **(Avalos et al., 2025)**. Consistent with a general shift toward more conservative strategies, height was decreased and diameter was increased. This suggests that warming induces stress requiring structural changes despite optimal water availability. These patterns align with the evidence that phenotypic plasticity to warming enables partial shifts along acquisitive–conservative continua **(Lohbeck et al., 2013)**.

#### 4.2.4. Warming reduces root investment under non-limiting water conditions

Warming reduced root-to-shoot in *Q. petraea,* whereas it increased for warmer *Q. pubescens* populations until a certain threshold, as expected for species inhabiting Mediterranean and submediterranean climates (Vicente et al., 2025). The observed decrease in root investment of *Q. petraea* populations under warmer-than-home conditions may reflect our experimental design, which ensured optimal water availability. This finding suggests that warming alone, without water limitation, does not stimulate greater root investment in this species. Consistent with this interpretation, identical results were found in *Q. robur*, *Q. petraea*, and *Q. pubescens*, where root:shoot ratio decreased with warming but increased with drought **(An et al., 2017; Arend et al., 2011)**.

The fine:main root ratio increased consistently under warming in both species, aligning with global patterns **(Wang et al., 2021)**. Under well-watered conditions, higher temperatures increase metabolic activity and atmospheric evaporative demand, potentially enhancing transpirational flux. This can elevate both water and nutrient demand. In response, seedlings may allocate more carbon to absorptive fine roots, which are primarily responsible for water and nutrient uptake, while investment in structural main roots decreases when soil water is not limiting **(Huang and Eissenstat, 2000; Wang et al., 2020)** but see **(Hertel et al., 2013)**.

## 5. Conclusion

Our findings demonstrate the importance of population climate in shaping germination and post-germination responses to warming in sessile and pubescent oak. The nature of this population effect is species-specific and reflects contrasting locally adapted strategies that generate unexpected vulnerabilities. In *Q. pubescens*, warmer and drier populations carry the cost of drought avoidance strategies under continuously wet conditions, performing worse than cooler populations despite their apparent pre-adaptation to hotter futures. In *Q. petraea*, continental populations, initially advantaged under moderate warming, suffer the steepest fitness declines under further warming, consistent with a narrow thermal optimum shaped by adaptation to cold.

Because these dynamics are highly species- and trait-specific, and because the responses observed here may be further modified under drought conditions (which are likely to alter greater root investment and fitness outcomes) our results underscore that regeneration potential under climate change cannot be generalized across species or populations. Only case-by-case assessments, integrating both climatic origin and early-stage trait responses to future conditions, can reliably inform assisted migration and regeneration strategies.

## Ackowledgments

We are grateful to Benjamin Dencausse and Aurélien Kohler for their exhaustive technical support during seedling phenotypic monitoring. We are also very thankful to Georgeta Mihai, Francisco Lario Leza, Maurizio Marchi, Giovanni Giuseppe Vendramin, Marcella Van Loo, Marjana Westergren, Bruno Fady, Thomas Sim, Johan Kroon, Jon Kehlet Hansen and Christian Rellstab for seed collection and shipment.

## Declarations

### Contributions by the Authors

MBG and MC designed the study. EV, MC and MBG conducted the sowing experiment and monitored germination. MC analyzed the data. MC and MBG wrote the manuscript. All the authors collected the acorns and edited the final manuscript. MBG acquired the financial support needed for this project.

