## Supplementary figures and tables for "Climate at seed origin drives germination and post-germination trait responses to warming in sessile and pubescent oaks"

### Supplementary material

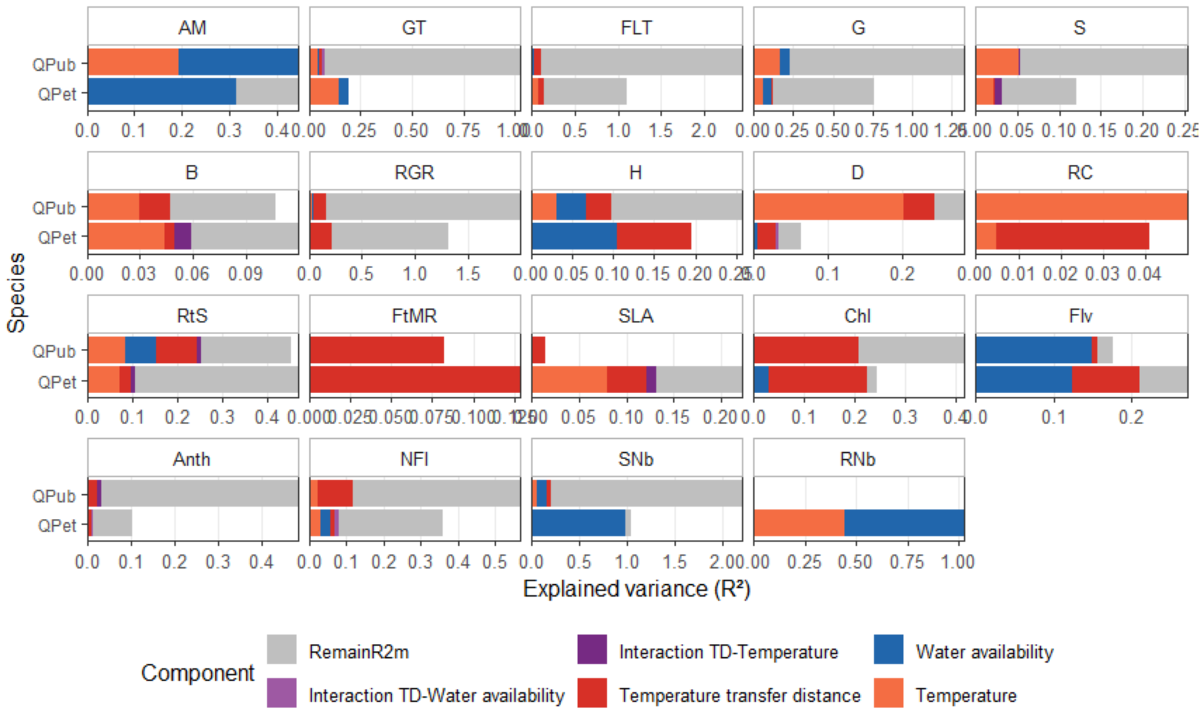


***Figure S1*** Explained variance partitioning for each generalized linear mixed-model of the effects of population and temperature transfer distance (including experiment temperature) on seed germination and seedling traits in *Q. pubescens* (QPub) and *Q. petraea* (QPet).

*In blue, population water availability variables explained part; in orange, population temperature variables; in red, temperature transfer distance capturing phenotypic plasticity and genetic variation in plasticity; in purple (light and dark), the interaction between transfer distance and population climate; in grey, the remaining part of marginal R-squared (partR2 quantifies semi-partial r-squared, representing the variance uniquely explained by each predictor; their sum is lower than the model's marginal R-squared because correlated predictors share explained variance that cannot be uniquely assigned to individual variables). The remaining part of conditional R-squared is not shown for the sake of readability. Traits acronyms mean: AM = Acorn Mass; GT = Germination Time; FLT = First Leaf Emergence time; G = Germination; S = Survival; B = total Biomass; RGR = Relative Growth Rate; H = Height; D = Diameter; RC = Root Collar; RtS = Root to Shoot ratio; FtMR = Fine to Main Roots ratio; SLA = Specific Leaf Area; Chl = Chlorophyll; Flv = Flavonoids; Anth = Anthocynanins; NFI = Nitrogen-Flavonoids Index; SNb = Stem Number; RNb = Root Number.*

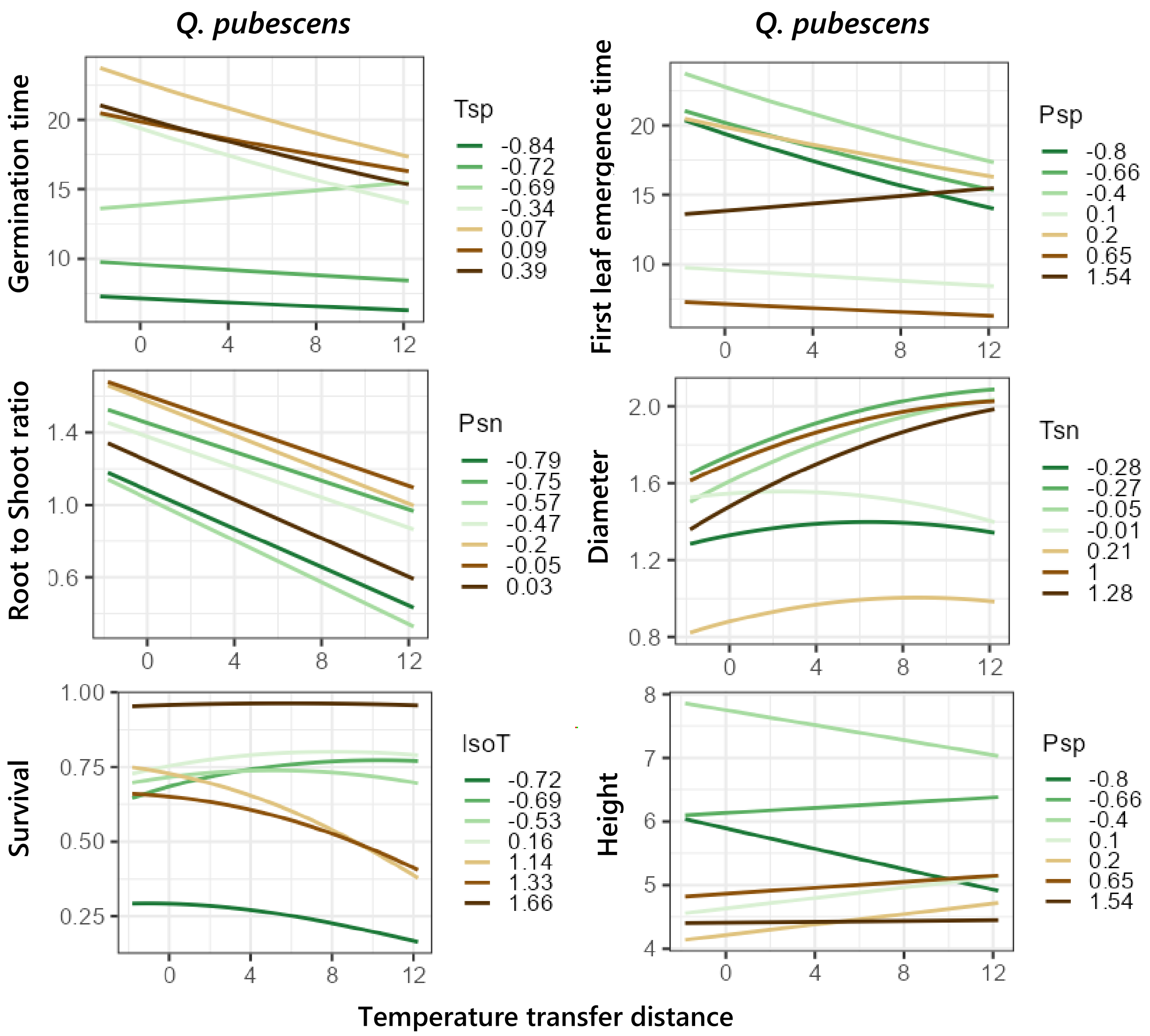


***Figure S2*** Plots showing the effects of temperature transfer distance and of the second most important population climate variable for each generalized linear mixed-model, for traits in which there was more than one significant population climate variable.

*Colored lines are populations represented function of the population climate variable that explained most variance. Climate variables acronyms mean: Tsp = mean spring Temperature; Psp = mean spring Precipitation; Psn = Precipitation seasonality; Tsn = Temperature seasonality; IsoT = Isothermality. No suffix indicates the variable is averaged on the 1901-1960 period. _AcMat suffix indicates the variable is averaged on the acorn maturation year.*

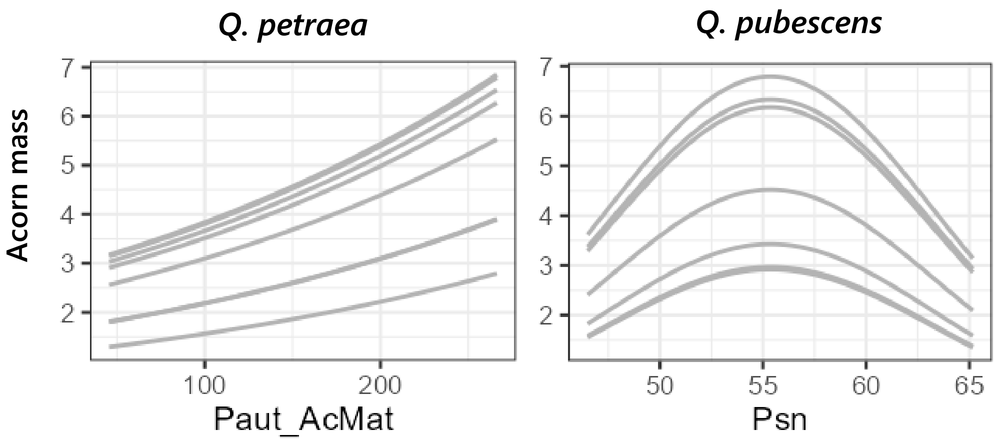


*Figure S3* Plots showing the effects of temperature transfer distance and of the most important population climate variable for each generalized linear mixed-model, for acorn mass.

*Lines are populations represented function of the population climate variable that explained most variance. Climate variables acronyms mean: Psn = Precipitation seasonality; Paut = mean autumn Precipitation. No suffix indicates the variable is averaged on the 1901-1960 period. _AcMat suffix indicates the variable is averaged on the acorn maturation year. Water availability during seed maturation controls maternal provisioning. In Q. petraea, acorn mass showed a positive relationship autumn precipitation during maturation. This positive? relationship is consistent in oaks (Koenig et al., 2013). Water stress in summer can interrupt acorn development even in Mediterranean oaks such as Q. ilex, leading to smaller size (Siscart et al., 1999; Alejano et al., 2011). Q pubescens acorns were bigger when growing at environments with intermediate precipitation seasonality This pattern may represent a reproductive allocation strategy balancing costs: under severe water stress (high seasonality), plants reduce investment in reproduction entirely; under abundant water (low seasonality), the need for large, resource-intensive acorns decreases; but under intermediate stress, producing larger seeds serves as a stress-coping strategy. This has been demonstrated in other species where populations under higher water stress produced larger seeds, including Q. ilex (Bonito et al., 2011), Q. suber (Ramirez-Valiente et al., 2009), and an Atlantic herbaceous species (Villellas & Garcia, 2012). Temperature during maturation showed weak effects on acorn mass compared to water availability, suggesting that water availability is the primary constraint on seed production in both species.*

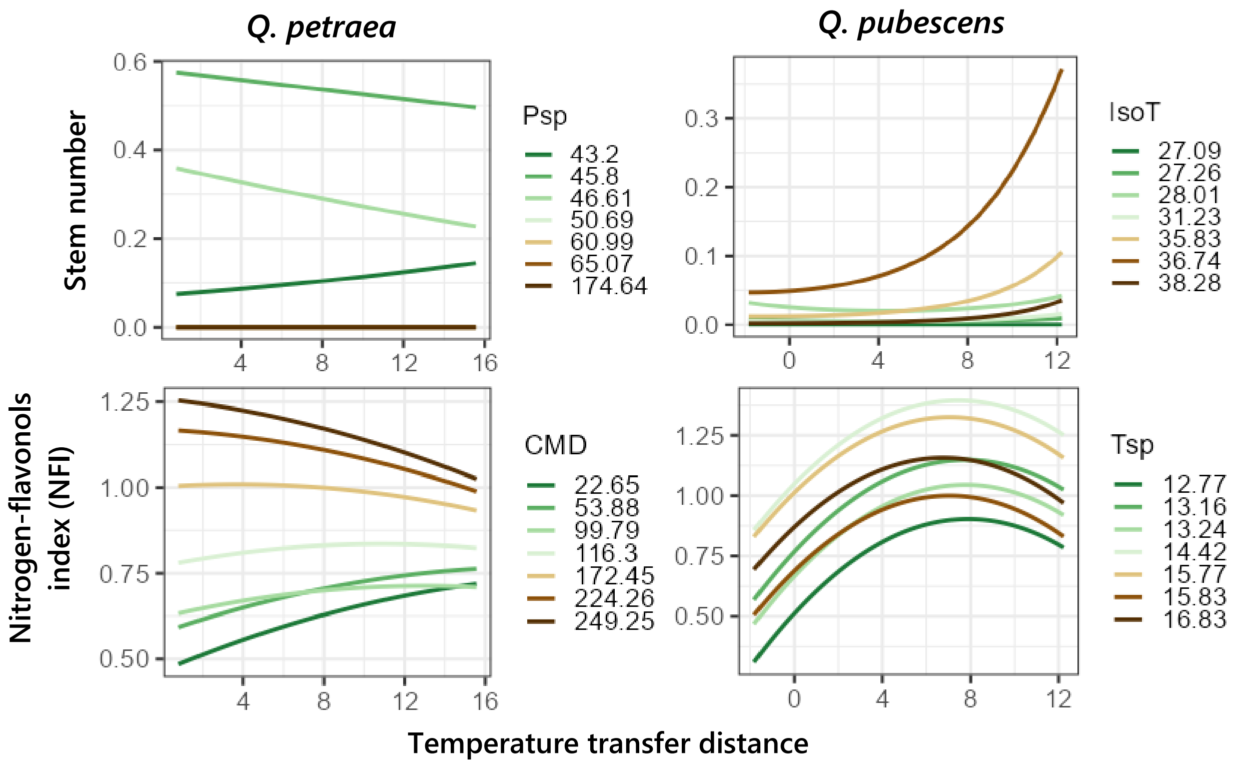


***Figure S4*** Plots showing the effects of temperature transfer distance and of the second most important population climate variable for each generalized linear mixed-model, for Stem number and NFI.

*Colored lines are populations represented function of the population climate variable that explained most variance. Climate variables acronyms mean: Tsp = mean spring Temperature; Psp = mean spring Precipitation; IsoT = Isothermality; CMD = Climatic Moisture Deficit Index. No suffix indicates the variable is averaged on the 1901-1960 period. _AcMat suffix indicates the variable is averaged on the acorn maturation year.*

| Method | R package | Reference |
|---|---|---|
| Stepwise selection | *caret 7.0-1* | **(Kuhn, 2008)** |
| Filter selection | *caret 7.0.1* | **(Kuhn, 2008)** |
| Random forest algorithm | *randomForest 4.7-1.2* | **(Liaw and Wiener, 2002)** |
| Boruta algorithm | *Boruta 9.0.0* | **(Kursa and Rudnicki, 2010)** |
| Recursive Feature Elimination | *caret 7.0.1* | **(Kuhn, 2008)** |
| Lasso Regularization | *glmnet 7.1-10* | **(Friedman et al., 2010)** |

***Table S1*** Multiple variable selection methods performed to build generalized linear mixed-effects model.

*We retained the variables that were selected the most, and the variables that had a higher importance rank in random forest and boruta methods, since these methods often show quite different output as they tend to capture nonlinear effects and interaction*

| Trait & general form | Formula | |
|---|---|---|
| | *Q. petraea* | *Q. pubescens* |
| AM $\sim \text{Gamma}(\mu, \phi)$ | $\log(\mu_{ij}) = \alpha_0 + \alpha_1 \text{Paut\_AcMat}_{ij} + \beta + \varepsilon$ | $\log(\mu_{ij}) = \alpha_0 + \alpha_1 \text{Tmin\_a\_AcMat}_{ij} + \alpha_2 \text{Tmin\_a\_AcMat}^2{}_{ij} + \alpha_3 \text{Psn}_{ij} + \alpha_4 \text{Psn}^2{}_{ij} + \beta + \varepsilon$ |
| G $\sim \text{Binomial}(n, p)$ | $\text{logit}(P(G_{ij} = 1)) = \alpha_0 + \alpha_1 \text{IsoT\_AcMat}_{ij} + \alpha_2 \text{Psn}_{ij} + \alpha_3 \text{TD}_{ijk} + \alpha_4 \text{Storage}_{ij} + \beta + \varepsilon$ | $\text{logit}(P(G_{ij} = 1)) = \alpha_0 + \alpha_1 \text{TD}_{ijk} + \alpha_2 \text{TD}^2{}_{ijk} + \alpha_3 \text{Tsp}_{ij} + \alpha_4 \text{Psn}_{ij} + \alpha_5 \text{TD}_{ijk} \times \text{Tsp}_{ij} + \beta + \varepsilon$ |
| S $\sim \text{Binomial}(n, p)$ | $\text{logit}(P(S_{ij} = 1)) = \alpha_0 + \alpha_1 \text{TD}_{ijk} + \alpha_2 \text{IsoT}_{ij} + \alpha_3 \text{TD}_{ijk} \times \text{IsoT}_{ij} + \beta + \varepsilon$ | $\text{logit}(P(S_{ij} = 1)) = \alpha_0 + \alpha_1 \text{TD}_{ijk} + \alpha_2 \text{TD}^2{}_{ijk} + \alpha_3 \text{Tmin\_a\_AcMat}_{ij} + \alpha_4 \text{IsoT}_{ij} + \alpha_5 \text{TD}_{ijk} \times \text{Tmin\_a\_AcMat}_{ij} + \beta + \varepsilon$ |
| GT $\sim \text{GenPoisson}(\mu, \phi)$ | $\log(\mu_{ij}) = \alpha_0 + \alpha_1 \text{TD}_{ij} + \alpha_2 \text{Psn\_AcMat}_{ij} + \alpha_3 \text{Tmin\_a\_AcMat}_{ij} + \alpha_4 \text{TD}_{ijk} \times \text{Psn\_AcMat}_{ij} + \beta + \varepsilon$ | $\log(\mu_{ij}) = \alpha_0 + \alpha_1 \text{TD}_{ijk} + \alpha_2 \text{Tsp}_{ij} + \alpha_3 \text{Tsp}^2{}_{ij} + \alpha_4 \text{Paut}_{ij} + \alpha_5 \text{TD}_{ijk} \times \text{Paut}_{ij} + \beta + \varepsilon$ |
| FLT $\sim \text{GenPoisson}(\mu, \phi)$ | $\log(\mu_{ij}) = \alpha_0 + \alpha_1 \text{TD}_{ijk} + \alpha_2 \text{Tsp}_{ij} + \alpha_3 \text{TD}_{ijk} \times \text{Tsp}_{ij} + \beta + \varepsilon$ | $\log(\mu_{ij}) = \alpha_0 + \alpha_1 \text{TD}_{ijk} + \alpha_2 \text{TD}^2{}_{ijk} + \alpha_3 \text{Tsp}_{ij} + \alpha_4 \text{Psp}_{ij} + \alpha_5 \text{Psp}^2{}_{ij} + \alpha_6 \text{TD}_{ijk} \times \text{Tsp}_{ij} + \beta + \varepsilon$ |
| B $\sim \mathcal{N}(\mu, \sigma^2)$ | $\mu_{ij} = \alpha_0 + \alpha_1 \text{TD}_{ij} + \alpha_2 \text{TD}^2{}_{ijk} + \alpha_3 \text{IsoT}_{ij} + \alpha_4 \text{TD}_{ijk} \times \text{IsoT}_{ij} + \beta + \varepsilon$ | $\mu_{ij} = \alpha_0 + \alpha_1 \text{TD}_{ijk} + \alpha_2 \text{Tsp}_{ij} + \alpha_3 \text{Psn}_{ij} + \beta + \varepsilon$ |
| RGR $\sim \text{Beta}(\mu, \phi)$ | $\text{logit}(\mu_{ij}) = \alpha_0 + \alpha_1 \text{TD}_{ijk} + \alpha_2 \text{TD}^2{}_{ijk} + \alpha_3 \text{CMD\_AcMat}^2{}_{ij} + \beta + \varepsilon$ | $\text{logit}(\mu_{ij}) = \alpha_0 + \alpha_1 \text{TD}_{ijk} + \alpha_2 \text{TD}^2{}_{ijk} + \alpha_3 \text{Psp}_{ij} + \alpha_4 \text{Tsp}_{ij} + \beta + \varepsilon$ |
| SLA $\sim \mathcal{N}(\mu, \sigma^2)$ | $\mu_{ij} = \alpha_0 + \alpha_1 \text{TD}_{ijk} + \alpha_2 \text{Tsn}_{ij} + \alpha_3 \text{Tsn}^2{}_{ij} + \alpha_4 \text{TD}_{ijk} \times \text{Tsn}_{ij} + \beta + \varepsilon$ | $\mu_{ij} = \alpha_0 + \alpha_1 \text{TD}_{ijk} + \beta + \varepsilon$ |
| Chl $\sim \mathcal{N}(\mu, \sigma^2)$ | $\mu_{ij} = \alpha_0 + \alpha_1 \text{TD}_{ijk} + \alpha_2 \text{TD}^2{}_{ijk} + \alpha_3 \text{CMD}_{ij} + \alpha_4 \text{TD}_{ijk} \times \text{Tsn}_{ij} + \beta + \varepsilon$ | $\mu_{ij} = \alpha_0 + \alpha_1 \text{TD}_{ijk} + \alpha_2 \text{TD}^2{}_{ijk} + \alpha_3 \text{Tsp}_{ij} + \alpha_4 \text{TD}_{ijk} \times \text{Tsp}_{ij} + \beta + \varepsilon$ |
| Flv $\sim \mathcal{N}(\mu, \sigma^2)$ | $\mu_{ij} = \alpha_0 + \alpha_1 \text{TD}_{ijk} + \alpha_2 \text{TD}^2{}_{ijk} + \alpha_3 \text{Psn\_AcMat}_{ij} + \alpha_4 \text{CMD}_{ij} + \beta + \varepsilon$ | $\mu_{ij} = \alpha_0 + \alpha_1 \text{TD}_{ijk} + \alpha_2 \text{TD}^2{}_{ijk} + \alpha_3 \text{Psp}_{ij} + \beta + \varepsilon$ |
| NFI $\sim \mathcal{N}(\mu, \sigma^2)$ | $\mu_{ij} = \alpha_0 + \alpha_1 \text{TD}_{ijk} + \alpha_2 \text{TD}^2{}_{ijk} + \alpha_3 \text{IsoT\_AcMat}_{ij} + \alpha_4 \text{IsoT\_AcMat}^2{}_{ij} + \alpha_5 \text{CMD}_{ij} + \alpha_5 (\text{TD}_{ijk} \times \text{CMD})_{ij} + \beta + \varepsilon$ | $\mu_{ij} = \alpha_0 + \alpha_1 \text{TD}_{ijk} + \alpha_2 \text{TD}^2{}_{ijk} + \alpha_3 \text{Tsp}_{ij} + \alpha_4 \text{Tsn}_{ij} + \alpha_5 \text{Tsn\_AcMat}_{ij} + \alpha_6 \text{TD}_{ijk} \times \text{Tsp}_{ij} + \beta + \varepsilon$ |

| Trait & general form | Formula | |
|---|---|---|
| | *Q. petraea* | *Q. pubescens* |
| Anth $\sim \mathcal{N}(\mu, \sigma^2)$ | $\mu_{ij} = \alpha_0 + \alpha_1 TD_{ijk} + \alpha_2 TD^2{}_{ijk} + \alpha_3 CMD_{ij} + \alpha_4 (TD_{ijk} \times CMD)_{ij} + \beta + \varepsilon$ | $\mu_{ij} = \alpha_0 + \alpha_1 TD_{ijk} + \alpha_2 TD^2{}_{ijk} + \alpha_3 Tsp_{ij} + \alpha_4 (TD_{ijk} \times Tsp)_{ij} + \beta + \varepsilon$ |
| H $\sim \mathcal{N}(\mu, \sigma^2)$ | $\mu_{ij} = \alpha_0 + \alpha_1 TD_{ijk} + \alpha_2 TD^2{}_{ijk} + \alpha_3 Psn\_AcMat_{ij} + \alpha_4 Psn\_AcMat^2{}_{ij} + \beta + \varepsilon$ | $\mu_{ij} = \alpha_0 + \alpha_1 TD_{ijk} + \alpha_2 TD^2{}_{ijk} + \alpha_3 Psp_{ij} + \alpha_4 Tsn_{ij} + \alpha_5 Tsn^2{}_{ij} + \alpha_6 (TD_{ijk} \times Tsn)_{ij} + \beta + \varepsilon$ |
| D $\sim \mathcal{N}(\mu, \sigma^2)$ | $\mu_{ij} = \alpha_0 + \alpha_1 TD_{ijk} + \alpha_2 TD^2{}_{ijk} + \alpha_3 Psn\_AcMat_{ij} + \alpha_4 Psn\_AcMat^2{}_{ij} + \alpha_5 (TD_{ijk} \times Psn\_AcMat)_{ij} + \beta + \varepsilon$ | $\mu_{ij} = \alpha_0 + \alpha_1 TD_{ijk} + \alpha_2 TD^2{}_{ijk} + \alpha_3 Tmin\_a\_AcMat_{ij} + \alpha_4 Tsn_{ij} + \alpha_5 (TD_{ijk} \times Tmin\_a\_AcMat)_{ij} + \beta + \varepsilon$ |
| RC $\sim \mathcal{N}(\mu, \sigma^2)$ | $\mu_{ij} = \alpha_0 + \alpha_1 TD_{ijk} + \alpha_2 TD^2{}_{ijk} + \alpha_3 IsoT_{ij} + \beta + \varepsilon$ | $\mu_{ij} = \alpha_0 + \alpha_1 Tmin\_a\_AcMat_{ij} + \beta + \varepsilon$ |
| RtS $\sim \mathcal{N}(\mu, \sigma^2)$ | $\mu_{ij} = \alpha_0 + \alpha_1 TD_{ijk} + \alpha_2 TD^2{}_{ijk} + \alpha_3 IsoT_{ij} + \alpha_4 IsoT^2{}_{ij} + \alpha_5 (TD_{ijk} \times IsoT)_{ij} + \beta + \varepsilon$ | $\mu_{ij} = \alpha_0 + \alpha_1 TD_{ijk} + \alpha_2 Tsp_{ij} + \alpha_3 Psn_{ij} + \alpha_4 (TD_{ijk} \times Tsp)_{ij} + \beta + \varepsilon$ |
| FtMR $\sim \mathcal{N}(\mu, \sigma^2)$ | $\mu_{ij} = \alpha_0 + \alpha_1 TD_{ijk} + \alpha_2 TD^2{}_{ijk} + \beta + \varepsilon$ | $\mu_{ij} = \alpha_0 + \alpha_1 TD_{ijk} + \alpha_2 TD^2{}_{ijk} + \beta + \varepsilon$ |
| SNb $\sim Binomial(n, p)$ | $\mu_{ij} = \alpha_0 + \alpha_1 TD_{ijk} + \alpha_2 Psp_{ij} + \alpha_3 (TD_{ijk} \times Psp)_{ij} + \beta + \varepsilon$ | $\mu_{ij} = \alpha_0 + \alpha_1 TD_{ijk} + \alpha_2 TD^2{}_{ijk} + \alpha_3 IsoT_{ij} + \alpha_4 Psp_{ij} + \alpha_5 Psp^2{}_{ij} + \alpha_6 (TD_{ijk} \times IsoT)_{ij} + \beta + \varepsilon$ |
| RNb $\sim Binomial(n, p)$ | $\mu_{ij} = \alpha_0 + \alpha_1 TD_{ijk} + \alpha_2 CMD_{ij} + \alpha_3 CMD^2{}_{ij} + \alpha_4 Tsp\_MTree_{ij} + \beta + \varepsilon$ | No significant relationship with the climatic variables tested |

***Table S2*** Description of the generalized mixed-models built per trait and species.

*The choice of the model type depended on the statistical distribution of each trait. In every equation, $ijk$ stands for the ith acorn of the jth population at the kth temperature. TD is the temperature transfer distance; $\alpha_n$ are the coefficients to be estimated; $Tsp$ = mean spring Temperature; $Tmin\_a$ = minimal annual Temperature; $Psn$ = Precipitation seasonality; $Paut$ = mean autumn precipitation; $IsoT$ = Isothermality; $Tsn$ = Temperature seasonality; $CMD$ = Climatic Moisture Deficit; $Psp$ = mean spring Precipitation; $\beta$ is the mother tree nested in the population as random effect; $\varepsilon$ the residuals. No suffix indicates the variable is averaged on the 1901-1960 period. $\_AcMat$ suffix indicates the variable is averaged on the acorn maturation year. In every model general form, $\mu$ is the mean parameter (function of predictor variables); $\phi$ is the precision parameter that controls the dispersion of the distribution; $\sigma^2$ is the variance of the Gaussian distribution; $n$ is the number of trials and $p$ is the probability of success (for the logistic regression). The mean parameter is linked to the regressors via a link function. Traits acronyms mean: $G$ = germination status ($G = 1$ means germinated; $G = 0$ means not germinated); $S$ = survival status ($S = 1$ means alive; $S = 0$ means dead); $AM$ = Acorn Mass; $GT$ = Germination*

*Time;* $FLT$ *= First Leaf Emergence time;* $B$ *= total Biomass;* $RGR$ *= Relative Growth Rate;* $H$ *= Height;* $D$ *= Diameter;* $RC$ *= Root Collar;* $RtS$ *= Root to Shoot ratio;* $FtMR$ *= Fine to Main Roots ratio;* $SLA$ *= Specific Leaf Area;* $Chl$ *= Chlorophyll;* $Flv$ *= Flavonoids;* $Anth$ *= Anthocynanins;* $NFI$ *= Nitrogen-Flavonoids Index;* $SNb$ *= Stem Number;* $RNb$ *= Root Number.*

| Species | Trait | AIC | AICc | BIC | $R^2$ cond. | $R^2$ marg. | RMSE | Rpearson_mean | Rpearson_SD |
|---|---|---|---|---|---|---|---|---|---|
| Q. petraea | GT | 20368,97 | 20369,02 | 20416,95 | 0,49 | 0,16 | 12,11 | 0,42 | 0,02 |
| Q. pubescens | | 16081,29 | 16081,36 | 16133,71 | 0,31 | 0,26 | 9,34 | 0,51 | 0,04 |
| Q. petraea | FLT | 12507,48 | 12507,57 | 12543,60 | 0,69 | 0,46 | 32,41 | 0,69 | 0,03 |
| Q. pubescens | | 13362,43 | 13362,58 | 13415,05 | 0,72 | 0,70 | 34,72 | 0,67 | 0,03 |
| Q. petraea | AM | 8360,18 | 8360,19 | 8390,87 | 0,69 | 0,27 | 0,82 | 0,81 | 0,01 |
| Q. pubescens | | 8041,98 | 8042,03 | 8089,52 | 0,63 | 0,44 | 1,01 | 0,73 | 0,02 |
| Q. petraea | G | 1881,19 | 1881,22 | 1923,98 | 0,47 | 0,28 | 0,28 | 0,43 | 0,05 |
| Q. pubescens | | 1537,17 | 1537,22 | 1584,71 | 0,63 | 0,45 | 0,28 | 0,44 | 0,04 |
| Q. petraea | S | 3606,77 | 3606,80 | 3642,75 | 0,22 | 0,06 | 0,44 | 0,31 | 0,04 |
| Q. pubescens | | 2785,60 | 2785,70 | 2832,20 | 0,28 | 0,10 | 0,42 | 0,35 | 0,04 |
| Q. petraea | RGR | -12195,40 | -12195,34 | -12156,77 | 1,00 | 0,77 | 0,02 | 0,33 | 0,04 |
| Q. pubescens | | -10053,31 | -10053,23 | -10009,99 | 1,00 | 0,78 | 0,02 | 0,43 | 0,04 |
| Q. petraea | B | 753,67 | 753,78 | 794,95 | 0,35 | 0,08 | 0,30 | 0,57 | 0,07 |
| Q. pubescens | | 1126,69 | 1126,77 | 1163,53 | 0,27 | 0,07 | 0,34 | 0,46 | 0,04 |
| Q. petraea | SLA | 14527,62 | 14527,71 | 14563,37 | 0,18 | 0,16 | 91,63 | 0,41 | 0,08 |
| Q. pubescens | | 14020,19 | 14020,24 | 14045,41 | 0,07 | 0,01 | 108,26 | 0,25 | 0,08 |
| Q. petraea | Chl | -1537,33 | -1537,23 | -1501,84 | 0,38 | 0,23 | 0,12 | 0,55 | 0,05 |
| Q. pubescens | | -805,20 | -805,07 | -765,17 | 0,38 | 0,28 | 0,16 | 0,57 | 0,05 |
| Q. petraea | Flv | -1424,59 | -1424,47 | -1384,03 | 0,42 | 0,23 | 0,12 | 0,58 | 0,04 |
| Q. pubescens | | -1279,17 | -1279,07 | -1244,15 | 0,41 | 0,17 | 0,13 | 0,56 | 0,04 |
| Q. petraea | NFI | 3137,02 | 3137,21 | 3187,72 | 0,26 | 0,15 | 0,37 | 0,31 | 0,08 |
| Q. pubescens | | 2730,51 | 2730,72 | 2780,55 | 0,26 | 0,21 | 0,32 | 0,42 | 0,05 |
| Q. petraea | Anth | -3480,80 | -3480,64 | -3435,19 | 0,24 | 0,05 | 0,08 | 0,28 | 0,09 |
| Q. pubescens | | -3018,89 | -3018,73 | -2973,86 | 0,24 | 0,18 | 0,09 | 0,45 | 0,11 |
| Q. petraea | H | 5579,48 | 5579,59 | 5620,76 | 0,35 | 0,18 | 1,99 | 0,46 | 0,04 |
| Q. pubescens | | 6360,91 | 6361,07 | 6413,53 | 0,24 | 0,14 | 2,15 | 0,44 | 0,05 |
| Q. petraea | D | 2347,52 | 2347,67 | 2393,51 | 0,20 | 0,04 | 0,59 | 0,36 | 0,06 |
| Q. pubescens | | 1694,16 | 1694,30 | 1740,91 | 0,40 | 0,25 | 0,43 | 0,52 | 0,02 |
| Q. petraea | RC | 3161,05 | 3161,15 | 3196,47 | 0,20 | 0,04 | 0,89 | 0,32 | 0,03 |
| Q. pubescens | | 2994,11 | 2994,16 | 3020,09 | 0,30 | 0,05 | 0,70 | 0,52 | 0,02 |
| Q. petraea | RtS | 3284,56 | 3284,70 | 3331,00 | 0,36 | 0,23 | 0,34 | 0,45 | 0,11 |
| Q. pubescens | | 4725,16 | 4725,26 | 4767,24 | 0,42 | 0,30 | 0,41 | 0,49 | 0,08 |
| Q. petraea | FtMR | -1911,44 | -1911,37 | -1880,96 | 0,28 | 0,13 | 0,10 | 0,46 | 0,04 |
| Q. pubescens | | -1661,35 | -1661,28 | -1630,14 | 0,31 | 0,08 | 0,11 | 0,35 | 0,07 |
| Q. petraea | SNb | 298,46 | 298,53 | 329,43 | 1,00 | 1,00 | 0,17 | 0,48 | 0,10 |
| Q. pubescens | | 457,32 | 457,43 | 506,06 | 0,76 | 0,70 | 0,18 | 0,32 | 0,09 |
| Q. petraea | RNb | 408,68 | 408,77 | 444,80 | 0,93 | 0,64 | 0,19 | 0,55 | 0,07 |

***Table S3*** Models performance measures.

*AIC the Akaike Information Criterion (measure of model quality that balances goodness-of-fit and model complexity); AICc the corrected Akaike Information Criterion (adjusted for small sample sizes); BIC the Bayesian Information Criterion (penalizes model complexity more strongly); $R^2$ cond. the conditional R-squared (proportion of variance explained by both fixed and random effects); $R^2$ marg. the marginal R-squared (proportion of variance explained by fixed effects); RMSE the Root Mean Square Error (average magnitude of model prediction errors); Rpearson_mean the Pearson correlation coefficient between observed and predicted values (averaged on 10 repetitions); Rpearson_SD the Pearson correlation coefficient standard deviation. Traits acronyms mean: AM = Acorn Mass; GT = Germination Time; FLT = First Leaf Emergence time; G = Germination; S = Survival; B = total Biomass; RGR = Relative Growth Rate; H = Height; D = Diameter; RC = Root Collar; RtS = Root to Shoot ratio; FtMR = Fine to Main Roots ratio; SLA = Specific Leaf Area; Chl = Chlorophyll; Flv = Flavonoids; Anth = Anthocynanins; NFI = Nitrogen-*

*Flavonoids Index; SNb = Stem Number; RNb = Root Number. Colors indicates how good marginal R-squared and R pearson are (from very low - red - to very high - blue).*

| Trait | Species | Covariate | semi-partial R2 | CI_lower | CI_upper | ndf |
|---|---|---|---|---|---|---|
| GT | *Q. petraea* | TD | 6,00E-04 | 0 | 0.208 | 4 |
| | | TD:Psn_AcMat | 0.0028 | 0 | 0.21 | 4 |
| | | Psn_AcMat | 0.0441 | 0 | 0.249 | 4 |
| | | Tmin_a_AcMat | 0.1462 | 0.0049 | 0.3453 | 4 |
| | *Q. pubescens* | I(Tsp^2) | 0 | 0 | 0.0279 | 5 |
| | | TD:Paut | 0.0105 | 0 | 0.0664 | 5 |
| | | TD | 0.0122 | 0 | 0.0681 | 5 |
| | | Paut | 0.0122 | 0 | 0.068 | 5 |
| | | Tsp | 0.0391 | 0 | 0.0941 | 5 |
| FLT | *Q. petraea* | TD:Tsp | 0.0034 | 0 | 0.0832 | 3 |
| | | TD | 0.0662 | 9,00E-04 | 0.1447 | 3 |
| | | Tsp | 0.0745 | 0.0029 | 0.1527 | 3 |
| | *Q. pubescens* | Psp | 4,00E-04 | 0 | 0.0584 | 6 |
| | | I(TD^2) | 0.0015 | 0 | 0.0594 | 6 |
| | | I(Psp^2) | 0.0066 | 5,00E-04 | 0.0642 | 6 |
| | | TD:Tsp | 0.0111 | 0.0016 | 0.0685 | 6 |
| | | Tsp | 0.0128 | 0.0021 | 0.0701 | 6 |
| | | TD | 0.085 | 0.0753 | 0.1384 | 6 |
| AM | *Q. petraea* | Paut_AcMat | 0.3142 | 0.1631 | 0.483 | 1 |
| | *Q. pubescens* | Tmin_a_AcMat | 0.0515 | 0.0708 | 0.2936 | 4 |
| | | I(Tmin_a_AcMat^2) | 0.1598 | 0.1814 | 0.3782 | 4 |
| | | I(Psn^2) | 0.25 | 0.2737 | 0.4487 | 4 |
| | | Psn | 0.251 | 0.2746 | 0.4494 | 4 |
| G | *Q. petraea* | TD | 0.0026 | 0 | 0.0832 | 4 |
| | | Storage | 0.0103 | 0 | 0.0911 | 4 |
| | | Psn | 0.0582 | 0.0124 | 0.1404 | 4 |
| | | IsoT_AcMat | 0.0596 | 0.0134 | 0.1419 | 4 |
| | *Q. pubescens* | TD:Tsp | 0 | 0 | 0.244 | 5 |
| | | TD | 3,00E-04 | 0 | 0.2442 | 5 |
| | | I(TD^2) | 0.0067 | 0 | 0.2507 | 5 |
| | | Psn | 0.0664 | 0 | 0.3108 | 5 |
| | | Tsp | 0.1562 | 0.0534 | 0.4013 | 5 |
| S | *Q. petraea* | TD | 0.0013 | 0 | 0.0181 | 3 |
| | | TD:IsoT | 0.008 | 0 | 0.0249 | 3 |
| | | IsoT | 0.0219 | 6,00E-04 | 0.0388 | 3 |
| | *Q. pubescens* | I(TD^2) | 4,00E-04 | 0 | 0.0394 | 5 |
| | | Tmin_a_AcMat | 7,00E-04 | 0 | 0.0397 | 5 |
| | | TD:Tmin_a_AcMat | 0.0011 | 0 | 0.04 | 5 |
| | | TD | 0.0068 | 0 | 0.0458 | 5 |

| Trait | Species | Covariate | semi-partial R2 | CI_lower | CI_upper | ndf |
|---|---|---|---|---|---|---|
| | | IsoT | 0.0193 | 0.0029 | 0.0585 | 5 |
| RGR | ***Q. petraea*** | CMD_AcMat | 0.003 | 0 | 0.0869 | 3 |
| | | I(TD^2) | 0.0069 | 0 | 0.0904 | 3 |
| | | TD | 0.0893 | 0.0692 | 0.1624 | 3 |
| | ***Q. pubescens*** | Psp | 0.0182 | 0 | 0.1008 | 4 |
| | | Tsp | 0.0291 | 0 | 0.1103 | 4 |
| | | I(TD^2) | 0.0686 | 0.0026 | 0.1447 | 4 |
| | | TD | 0.1192 | 0.0137 | 0.1887 | 4 |
| B | ***Q. petraea*** | I(TD^2) | 0.0066 | 0 | 0.0731 | 4 |
| | | TD | 0.0069 | 0 | 0.0734 | 4 |
| | | TD:IsoT | 0.0101 | 0 | 0.0764 | 4 |
| | | IsoT | 0.0443 | 0.0024 | 0.1089 | 4 |
| | ***Q. pubescens*** | Psn | 0 | 0 | 0.1361 | 3 |
| | | TD | 0.0168 | 0 | 0.155 | 3 |
| | | Tsp | 0.03 | 0.002 | 0.167 | 3 |
| SLA | ***Q. petraea*** | Tsn:TD | 0.0113 | 0 | 0.1063 | 4 |
| | | I(Tsn^2) | 0.0266 | 0 | 0.1197 | 4 |
| | | TD | 0.0408 | 0 | 0.1323 | 4 |
| | | Tsn | 0.0694 | 0.0028 | 0.1574 | 4 |
| | ***Q. pubescens*** | TD | 0.0145 | 0.0073 | 0.0257 | 1 |
| Chl | ***Q. petraea*** | I(TD^2) | 0.0232 | 0 | 0.1221 | 3 |
| | | CMD | 0.0309 | 0 | 0.1287 | 3 |
| | | TD | 0.1125 | 0.0292 | 0.1983 | 3 |
| | ***Q. pubescens*** | Tsp | 0 | 0 | 0.026 | 4 |
| | | TD:Tsp | 4,00E-04 | 0 | 0.0573 | 4 |
| | | I(TD^2) | 0.042 | 0.0013 | 0.0969 | 4 |
| | | TD | 0.1952 | 0.1695 | 0.2427 | 4 |
| Flv | ***Q. petraea*** | Psn_AcMat | 0 | 0 | 0.1813 | 4 |
| | | I(TD^2) | 6,00E-04 | 0 | 0.1824 | 4 |
| | | TD | 0.0307 | 0 | 0.2081 | 4 |
| | | CMD | 0.1238 | 0.0014 | 0.2877 | 4 |
| | ***Q. pubescens*** | TD | 0 | 0 | 0.1071 | 3 |
| | | I(TD^2) | 0.0043 | 0 | 0.1128 | 3 |
| | | Psp | 0.1496 | 0.03 | 0.2401 | 3 |
| NFI | ***Q. petraea*** | I(TD^2) | 0.0013 | 0 | 0.2118 | 6 |
| | | TD | 0.0061 | 0 | 0.2158 | 6 |
| | | I(IsoT_AcMat^2) | 0.0075 | 0 | 0.2168 | 6 |
| | | TD:CMD | 0.0125 | 0 | 0.221 | 6 |

| Trait | Species | Covariate | semi-partial R2 | CI_lower | CI_upper | ndf |
|---|---|---|---|---|---|---|
| | | IsoT_AcMat | 0.0138 | 0 | 0.222 | 6 |
| | | CMD | 0.0291 | 0 | 0.2346 | 6 |
| | *Q. pubescens* | Tsn_AcMat | 0 | 0 | 0.0818 | 6 |
| | | TD:Tsp | 5,00E-04 | 0 | 0.0824 | 6 |
| | | Tsn | 0.0069 | 0 | 0.0879 | 6 |
| | | Tsp | 0.017 | 0 | 0.0967 | 6 |
| | | I(TD^2) | 0.03 | 0 | 0.108 | 6 |
| | | TD | 0.0771 | 0.0419 | 0.149 | 6 |
| Anth | *Q. petraea* | I(TD^2) | 0.001 | 0 | 0.0417 | 4 |
| | | TD | 0.0036 | 0 | 0.0442 | 4 |
| | | TD:CMD | 0.0036 | 0 | 0.0442 | 4 |
| | | CMD | 0.0049 | 1,00E-04 | 0.0454 | 4 |
| | *Q. pubescens* | Tsp | 0 | 0 | 0.1571 | 4 |
| | | TD:Tsp | 0.0095 | 0.0049 | 0.1761 | 4 |
| | | TD | 0.0117 | 0.0076 | 0.1783 | 4 |
| | | I(TD^2) | 0.016 | 0.0124 | 0.1828 | 4 |
| H | *Q. petraea* | I(TD^2) | 0.0051 | 0 | 0.0742 | 4 |
| | | TD | 0.0428 | 0.0018 | 0.1091 | 4 |
| | | I(Psn_AcMat^2) | 0.1024 | 0.0613 | 0.1643 | 4 |
| | | Psn_AcMat | 0.107 | 0.0661 | 0.1686 | 4 |
| | *Q. pubescens* | TD:Tsn | 0 | 0 | 0.0565 | 6 |
| | | I(Tsn^2) | 0.0201 | 0.0039 | 0.075 | 6 |
| | | TD | 0.0242 | 0.0079 | 0.0786 | 6 |
| | | Tsn | 0.0289 | 0.0127 | 0.0829 | 6 |
| | | I(TD^2) | 0.0301 | 0.0139 | 0.084 | 6 |
| | | Psp | 0.0347 | 0.0185 | 0.0882 | 6 |
| D | *Q. petraea* | TD:Psn_AcMat | 0.004 | 0 | 0.0728 | 5 |
| | | Psn_AcMat | 0.0136 | 0 | 0.0821 | 5 |
| | | I(Psn_AcMat^2) | 0.0142 | 0 | 0.0827 | 5 |
| | | I(TD^2) | 0.017 | 6,00E-04 | 0.0854 | 5 |
| | | TD | 0.0238 | 0.0045 | 0.092 | 5 |
| | *Q. pubescens* | I(TD^2) | 0 | 0 | 0.1425 | 5 |
| | | TD:Tmin_a_AcMat | 0 | 0 | 0.1285 | 5 |
| | | TD | 0.0195 | 0.001 | 0.159 | 5 |
| | | Tsn | 0.0567 | 0.0092 | 0.1897 | 5 |
| | | Tmin_a_AcMat | 0.1447 | 0.0962 | 0.2621 | 5 |
| RC | *Q. petraea* | IsoT | 0.0049 | 0 | 0.0291 | 3 |
| | | I(TD^2) | 0.0332 | 0.0227 | 0.0574 | 3 |
| | | TD | 0.0345 | 0.024 | 0.0588 | 3 |

| Trait | Species | Covariate | semi-partial R2 | CI_lower | CI_upper | ndf |
|---|---|---|---|---|---|---|
| | ***Q. pubescens*** | Tmin_a_AcMat | 0.0497 | 0.0018 | 0.1202 | 1 |
| **RtS** | ***Q. petraea*** | IsoT | 2,00E-04 | 0 | 0.0993 | 5 |
| | | TD | 0.0042 | 0 | 0.1029 | 5 |
| | | TD:IsoT | 0.0102 | 0 | 0.1081 | 5 |
| | | I(TD^2) | 0.0229 | 0 | 0.1192 | 5 |
| | | I(IsoT^2) | 0.0634 | 0.0304 | 0.1548 | 5 |
| | ***Q. pubescens*** | TD:Tsp | 0.0076 | 0 | 0.1589 | 4 |
| | | Psn | 0.0696 | 0 | 0.2102 | 4 |
| | | Tsp | 0.0846 | 0 | 0.2226 | 4 |
| | | TD | 0.091 | 0 | 0.2279 | 4 |
| **FtMR** | ***Q. petraea*** | I(TD^2) | 0 | 0 | 0.0199 | 2 |
| | | TD | 0.0285 | 4,00E-04 | 0.0466 | 2 |
| | ***Q. pubescens*** | I(TD^2) | 0 | 0 | 0.0204 | 2 |
| | | TD | 0.0387 | 0.0225 | 0.0614 | 2 |
| **RNb** | ***Q. petraea*** | TD | 0 | 0 | 0.9129 | 4 |
| | | Tsp | 0.3453 | 0 | 0.9499 | 4 |
| | | I(CMD^2) | 0.5028 | 0.1283 | 0.9608 | 4 |
| | | CMD | 0.5041 | 0.1296 | 0.9609 | 4 |
| **SNb** | ***Q. petraea*** | TD | 0 | 0 | 0.7463 | 3 |
| | | TD:Psp | 0 | 0 | 0.7458 | 3 |
| | | Psp | 0.9828 | 0.8234 | 0.9981 | 3 |
| | ***Q. pubescens*** | Psp | 0 | 0 | 0.1116 | 6 |
| | | TD:IsoT | 0 | 0 | 0.0942 | 6 |
| | | I(TD^2) | 0.006 | 0 | 0.1245 | 6 |
| | | TD | 0.0082 | 0 | 0.1262 | 6 |
| | | IsoT | 0.0625 | 0 | 0.1671 | 6 |
| | | I(Psp^2) | 0.1019 | 0.0079 | 0.1987 | 6 |

***Table S4*** Semi-partial $R^2$ values for fixed effects estimated using the partR2 R package. *Semi-partial $R^2$ represents the proportion of variance in the response uniquely explained by each covariate after accounting for all other fixed effects in the model. Confidence intervals (CI_lower, CI_upper) were obtained by parametric bootstrapping. ndf indicates the number of degrees of freedom associated with each covariate. Within each group, semi-partial $R^2$ are sorted in ascending order.*

| Trait | Species | Covariate | Estimate | p |
|---|---|---|---|---|
| AM | QPet | **Paut_AcMat** | **0.2** | **0.001** |
| | QPub | Tmin_a_AcMat | 0.11 | 0.4 |
| | | **Tmin_a_AcMat²** | **-1.37** | **0.001** |
| | | **Psn** | **-3.33** | **0.001** |
| | | **Psn²** | **-4.14** | **0.001** |
| G | QPet | **IsoT_AcMat** | **-2.57** | **0.001** |
| | | **Psn** | **-2.84** | **0.001** |
| | | **TD** | **-0.15** | **0.01** |
| | | Storage | -0.31 | 0.44 |
| | QPub | TD | 0.01 | 0.91 |
| | | **TD²** | **-0.35** | **0.001** |
| | | **Tsp** | **-2.98** | **0.001** |
| | | Psn | 1.91 | 0.1 |
| | | **TD x Tsp** | **-0.96** | **0.001** |
| GT | QPet | **TD** | **-0.05** | **0.05** |
| | | Psn_AcMat | -1.03 | 0.12 |
| | | **Tmin_a_AcMat** | **-1.01** | **0.02** |
| | | **TD x Psn_AcMat** | **-0.09** | **0.02** |
| | QPub | **TD** | **-0.06** | **0.001** |
| | | Tsp | 0.32 | 0.06 |
| | | **Tsp²** | **-1.09** | **0.001** |
| | | **Paut** | **0.12** | **0.001** |
| | | **TD x Paut** | **0.05** | **0.001** |
| FLT | QPet | **TD** | **-0.34** | **0.001** |
| | | **Tsp** | **-0.46** | **0.001** |
| | | **TD x Tsp** | **-0.07** | **0.001** |
| | QPub | **TD** | **-0.34** | **0.001** |
| | | **TD²** | **0.04** | **0.01** |
| | | **Tsp** | **-0.94** | **0.001** |
| | | Psp | -0.08 | 0.09 |
| | | **Psp²** | **-0.09** | **0.03** |
| | | **TD x Tsp** | **0.2** | **0.001** |
| S | QPet | **TD** | **-0.11** | **0.03** |
| | | **IsoT** | **-0.82** | **0.001** |
| | | **TD x IsoT** | **0.22** | **0.001** |
| | QPub | TD | 0.05 | 0.48 |
| | | TD² | -0.06 | 0.17 |
| | | **Tmin_a_AcMat** | **-1.21** | **0.001** |
| | | **IsoT** | **0.3** | **0.05** |
| | | **TD x Tmin_a_AcMat** | **-0.47** | **0.001** |
| RGR | QPet | **TD** | **-0.38** | **0.001** |
| | | TD² | 0.04 | 0.06 |

| Trait | Species | Covariate | Estimate | p |
|---|---|---|---|---|
| | | CMD_AcMat | 0.27 | 0.1 |
| | QPub | **TD** | **-0.38** | **0.001** |
| | | **$TD^2$** | **0.16** | **0.001** |
| | | **Psp** | **-0.29** | **0.001** |
| | | Tsp | 0.17 | 0.25 |
| B | QPet | TD | -0.01 | 0.54 |
| | | **$TD^2$** | **-0.05** | **0.001** |
| | | IsoT | **-0.2** | **0.02** |
| | | **TD x IsoT** | **0.05** | **0.001** |
| | QPub | **TD** | **-0.05** | **0.001** |
| | | **Tsp** | **-0.28** | **0.02** |
| | | Psn | -0.1 | 0.53 |
| SLA | QPet | **Tsn** | **52.39** | **0.001** |
| | | **TD** | **-22.07** | **0.001** |
| | | $Tsn^2$ | 9.43 | 0.27 |
| | | **TD x Tsn** | **-17.64** | **0.001** |
| | QPub | **TD** | **-12.01** | **0.001** |
| Chl | QPet | **CMD** | 0.07 | 0.14 |
| | | **TD** | **0.09** | **0.001** |
| | | **$TD^2$** | **-0.01** | **0.001** |
| | QPub | **TD** | **0.13** | **0.001** |
| | | **$TD^2$** | **-0.06** | **0.001** |
| | | **Tsp** | **0.21** | **0.001** |
| | | **TD x Tsp** | **-0.06** | **0.001** |
| Flv | QPet | **TD** | **0.05** | **0.001** |
| | | $TD^2$ | 0 | 0.42 |
| | | Psn_AcMat | 0 | 0.95 |
| | | **CMD** | **-0.2** | **0.01** |
| | QPub | **Psp** | **0.1** | **0.001** |
| | | TD | -0.01 | 0.46 |
| | | **$TD^2$** | **0.01** | **0.01** |
| Anth | QPet | **TD** | **0.26** | **0.001** |
| | | $TD^2$ | 0.01 | 0.49 |
| | | CMD | 0.37 | 0.18 |
| | | **TD x CMD** | **0.28** | **0.001** |
| | QPub | **TD** | **0.24** | **0.001** |
| | | **$TD^2$** | **-0.19** | **0.001** |
| | | **Tsp** | **0.75** | **0.001** |
| | | **TD x Tsp** | **-0.4** | **0.001** |
| NFI | QPet | **TD** | **-0.11** | **0.001** |
| | | $TD^2$ | 0 | 0.64 |
| | | IsoT_AcMat | -0.02 | 0.91 |

| Trait | Species | Covariate | Estimate | p |
|---|---|---|---|---|
| | | IsoT_AcMat$^2$ | 0.23 | 0.65 |
| | | **CMD** | **0.65** | **0.001** |
| | | **TD x CMD** | **-0.14** | **0.001** |
| | QPub | **TD** | **0.13** | **0.001** |
| | | **TD$^2$** | **-0.1** | **0.001** |
| | | **Tsp** | **0.36** | **0.001** |
| | | Tsn | 0.44 | 0.06 |
| | | Tsn_AcMat | -0.37 | 0.15 |
| | | **TD x Tsp** | **-0.13** | **0.001** |
| RC | QPet | **TD** | **0.36** | **0.001** |
| | | **TD$^2$** | **-0.15** | **0.001** |
| | | IsoT | -0.17 | 0.13 |
| | QPub | **Tmin_a_AcMat** | **-0.47** | **0.03** |
| RtS | QPet | TD | 0.04 | 0.1 |
| | | **TD$^2$** | **-0.04** | **0.001** |
| | | IsoT | 0 | 1 |
| | | **IsoT$^2$** | **0.19** | **0.001** |
| | | **TD x IsoT** | **0.09** | **0.001** |
| | QPub | **TD** | **-0.24** | **0.001** |
| | | **Tsp** | **-0.58** | **0.001** |
| | | **Psn** | **0.45** | **0.01** |
| | | **TD x Tsp** | **-0.1** | **0.001** |
| FtMR | QPet | **TD** | **0.03** | **0.001** |
| | | TD$^2$ | 0 | 0.18 |
| | QPub | **TD** | **0.03** | **0.001** |
| | | TD$^2$ | 0 | 0.44 |
| H | QPet | **TD** | **-0.86** | **0.001** |
| | | **TD$^2$** | **0.14** | **0.001** |
| | | **Psn_AcMat** | **-12.42** | **0.001** |
| | | **Psn_AcMat$^2$** | **-11.43** | **0.001** |
| | QPub | **Psp** | **-1.04** | **0.03** |
| | | **TD** | **-0.51** | **0.001** |
| | | **TD$^2$** | **0.36** | **0.001** |
| | | Tsn | 2.37 | 0.08 |
| | | Tsn$^2$ | -2.06 | 0.18 |
| | | **TD x Tsn** | **-0.24** | **0.01** |
| D | QPet | **TD** | **0.31** | **0.001** |
| | | **TD$^2$** | **-0.07** | **0.001** |
| | | Psn_AcMat | -1.34 | 0.14 |
| | | Psn_AcMat$^2$ | -1.01 | 0.22 |
| | | **TD x Psn_AcMat** | **0.17** | **0.01** |
| | QPub | **TD** | **0.13** | **0.001** |

| Trait | Species | Covariate | Estimate | p |
|---|---|---|---|---|
| | | **TD²** | **-0.02** | **0.04** |
| | | **Tmin_a_AcMat** | **-0.47** | **0.001** |
| | | **Tsn** | **-0.15** | **0.04** |
| | | **TD x Tmin_a_AcMat** | **-0.16** | **0.001** |
| SNb | QPet | TD | -2.89 | 0.38 |
| | | Psp | -38.6 | 0.08 |
| | | TD x Psp | -4.07 | 0.38 |
| | QPub | **TD** | **0.4** | **0.01** |
| | | **TD²** | **1.21** | **0.001** |
| | | IsoT | -0.56 | 0.5 |
| | | Psp | 0.43 | 0.54 |
| | | Psp² | -3.96 | 0.17 |
| | | **TD x IsoT** | **0.34** | **0.02** |
| RNb | QPet | TD | -0.02 | 0.87 |
| | | CMD | -95.8 | 0.35 |
| | | CMD² | -47.54 | 0.34 |
| | | Tsp_MTree | 1.9 | 0.64 |
| MC | QPet | **Psn** | **-0.23** | **0.001** |
| | | **Tsn_AcMat** | **-0.16** | **0.001** |
| | | **Tsn_AcMat²** | **-0.12** | **0.001** |
| | QPub | **Tmin_a_AcMat** | **-0.38** | **0.01** |
| | | **Psn** | **-0.42** | **0.03** |

***Table S5*** Parameters of each model

*Acorn Mass (AM), Germination (G), Germination Time (GT), and First Leaf emergence Time (FLT), Survival (S), Relative Growth Rate (RGR), Total Biomass (B), Chlorophyll (Chl), Flavonols (Flv), Anthocyanins (Anth), Nitrogen-Flavonols Index (NFI) and Surface Leaf Area Index (SLA), Height (H), Diameter (D), Stem Number (SNb), Root Collar (RC), Root to Shoot ratio (RtS), Fine roots to Main Roots ratio (FtMR), and main Root Number (RNb); for Q. petraea and Q. pubescens. Estimate are the coefficients of regression; p the p value. Significant p values ($p < 0.05$) are shown in bold. IsoT the temperature Isothermality; Paut the mean autumn precipitation; Psn the precipitation seasonality; TD the temperature transfer distance; Psp the mean spring precipitation; Tmin_a the annual minimal temperature; Tsp the mean spring temperature; CMD the Climatic Moisture Deficit index. No suffix added to the variable means it is averaged on 1901-1960; AcMat suffix means Acorn Maturation year. For better readability, variables are grouped with colors: population temperature variables are in orange; population water availability variables are in blue; temperature transfer distance is in red. Negative effects' cells are colored in light red; positive effects' cells are colored in light blue.*
